# The S-lignin *O*-demethylase SyoA: Structural insights into a new class of heme peroxygenase enzymes

**DOI:** 10.1101/2024.07.14.603228

**Authors:** Alix C. Harlington, Tuhin Das, Keith E. Shearwin, Stephen G. Bell, Fiona Whelan

## Abstract

The *O*-demethylation of lignin aromatics is a rate-limiting step in their bioconversion to high-value compounds. A recently discovered cytochrome P450 enzyme SyoA was found to demethylate the sinapyl alcohol-derived (S-lignin) aromatic syringol. In this work, we solved high-resolution X-ray crystal structures of SyoA in the substrate-free and substrate-bound states and evaluate the demethylation of *para*-substituted S-lignin aromatics via the monooxygenase pathway and peroxide shunt pathway. We found that SyoA demethylates S-lignin aromatics with the following activity: 4-methylsyringol > syringaldehyde > syringol exclusively using the peroxide shunt pathway. The atomic-resolution structure of SyoA reveals the position of the non-canonical residues in the I-helix (Gln252 and Glu253). Site-directed mutagenesis of this amide-acid pair of a homologous CYP255 enzyme GcoA, which can catalyze the O-demethylation of guaiacol using both monooxygenase and peroxygenase activity, showed the amide-acid pair is critical for both pathways. This work expands the enzymatic toolkit for improving the capacity to funnel lignin towards high-value compounds, and defines the new chemistry within the active site of the enzyme that enables efficient peroxygenase activity. These insights provide a framework for engineered peroxygenase activity in other cytochrome P450 enzymes, with the potential for more facile catalysis, relative to traditional P450 monooxygenases which require difficult to handle redox partners and expensive nicotinamide cofactors.

## Introduction

Ubiquitous in all domains of life, the superfamily of heme-dependent cytochrome P450 enzymes (P450s) catalyze selective oxidations of unactivated C−H bonds. In recent years, P450s have gained traction as catalysts for lignin valorization (1). The CYP255 family of P450s; so far consisting of GcoA (2), AgcA (3) and SyoA (4), catalyze the *O*-demethylation of lignin aromatics. Lignin is predominantly comprised of coniferyl (G-type) and sinapyl (S-type) alcohol subunits, which have one or two methoxy groups, respectively (5). Aromatics derived from lignin depolymerization must be *O*-demethylated to diols to permit formation of ring-opened compounds, which can be funneled to high-value compounds (6). So far, three classes of enzymes have been identified that catalyze *O*-demethylation of S-lignin aromatics including: tetrahydrofolate (THF)-dependent demethylases (7, 8); Rieske non-heme iron-dependent oxygenases (9, 10); and P450s (4) (Fig. 1A). P450 enzymes have been engineered to demethylate syringol, including GcoA (11) and CYP102A1 (P450_BM3_) (12); however, native enzymes capable of the *O*-demethylation of S-lignin aromatics remain largely undescribed.

**Fig. 1.**
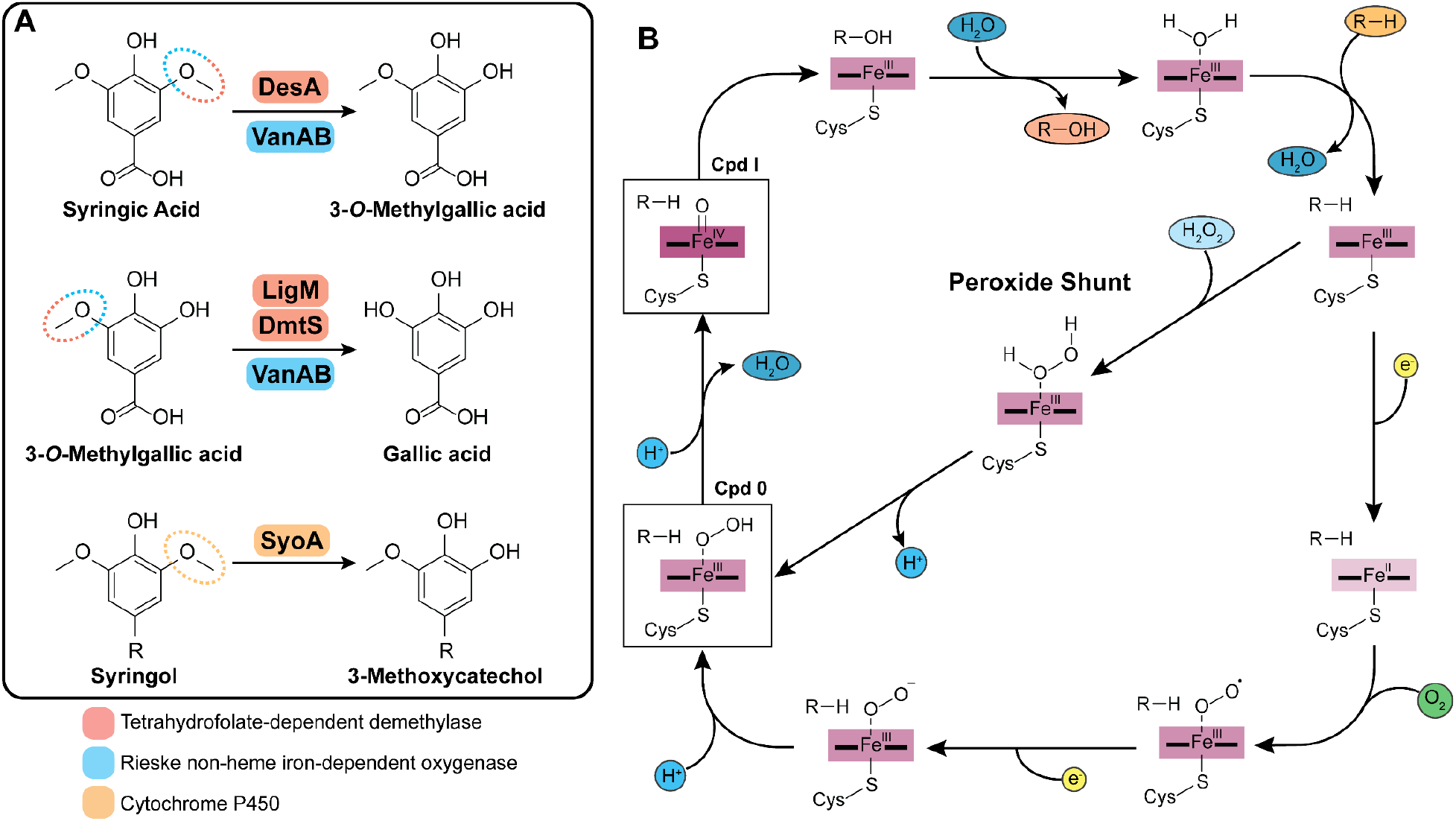
(A) Schemes showing the *O*-demethylation reactions of S-lignin aromatics catalyzed by tetrahydrofolate-dependent demethylases DesA, LigM and DmtS from *Sphingobium* sp. SYK-6 and *Novosphingobium aromaticivorans* DSM 12444, the Rieske non-heme iron-dependent oxygenase VanAB from *Pseudomonas putida* KT2440 and the cytochrome P450 SyoA from *Amycolatopsis thermoflava* N1165 (R = H, CH_3_, CHO). (B) Catalytic cycle of cytochrome P450s showing the monooxygenase pathway (outer circle) and the peroxide shunt pathway.

Most P450s function as monooxygenases requiring redox partners to transfer two electrons to the heme iron from the nicotinamide cofactor NAD(P)H to activate dioxygen (O_2_) (13, 14). Activation of dioxygen results in the formation of the oxy-ferryl porphyrin radical cation species compound I (Cpd I), the reactive intermediate (15, 16) (Fig. 1B). In many P450s this process is mediated by two residues within the oxygen binding groove of the I-helix—referred to as the acid-alcohol pair (Asp/Glu-Thr/Ser). Specifically, the acid aids in the delivery of protons to the heme-oxygen intermediates, while the alcohol is believed to stabilize the ferric-hydroperoxo intermediate, compound 0 (Cpd 0) for efficient formation of Cpd I (17– 21). P450s can also use H_2_O_2_ to generate Cpd I by using the ‘peroxide shunt pathway’, circumventing the need for additional redox partner proteins and expensive NAD(P)H cofactor (22, 23) (Fig. 1B). The majority of P450s are unable to use H_2_O_2_ efficiently, leading to heme destruction and enzyme inactivation (24). However, some families of heme enzymes possess naturally high levels of peroxygenase activity and deploy residues capable of acid-base chemistry to activate H_2_O_2_. For example, the CYP152 family use the substrate’s carboxyl group, coordinated to an Arg, to function as peroxygenases (25– 27). Furthermore, the CYP177 enzyme (*Ssca*CYP) was shown to hydroxylate *trans*-β-methyl-styrene and catalyze the sulfoxidation of thioanisole using H_2_O_2_ (28). Structural analysis of *Ssca*CYP revealed two acid-base residues in the active site; an Asp which replaces the canonical I-helix Thr, and a Glu near the K-helix. Other heme-containing peroxidases and peroxygenases also deploy residues capable of acid-base chemistry on the distal side of the heme (22, 29). It is believed that binding of H_2_O_2_ to the ferric form of these enzymes results in deprotonation of the iron-bound oxygen, forming Cpd 0, with a protonation step resulting in O–O bond scission and dehydration leading to Cpd I (Fig. 1C).

Here we report high-resolution X-ray crystal structures of the newly characterized CYP255 enzyme SyoA in the open substrate-free and closed substrate-bound states. These structures collectively revealed capacity in the active site to accommodate *O*-demethylation of *para*-substituted S-lignin aromatics. We demonstrate that SyoA functions more efficiently using the peroxide shunt pathway to demethylate S-lignin aromatics compared to the monooxygenase pathway. Additionally, we investigate the role of the amide-acid pair (QE) of residues, which sit above the heme within the I-helix, in the monooxygenase and peroxygenase activity of the CYP255 using GcoA as a model system. The activity of GcoA QE mutants was interrogated, illustrating that, for this residue pair, both monooxygenase and peroxygenase activities are profoundly impacted by reversion to the canonical P450 ET sequence.

## Results

### Structural analysis and conformational change of SyoA

To investigate syringol binding by SyoA, we performed structural characterization by X-ray crystallography. Crystal structures of substrate-free (PDB: 8u09) and syringol-bound (PDB: 8u19) SyoA were determined at resolutions of 1.98 and 1.26 Å, respectively (Fig. 2; SI Appendix, Fig. S1, S2 A and B and Table S1).

**Fig. 2.**
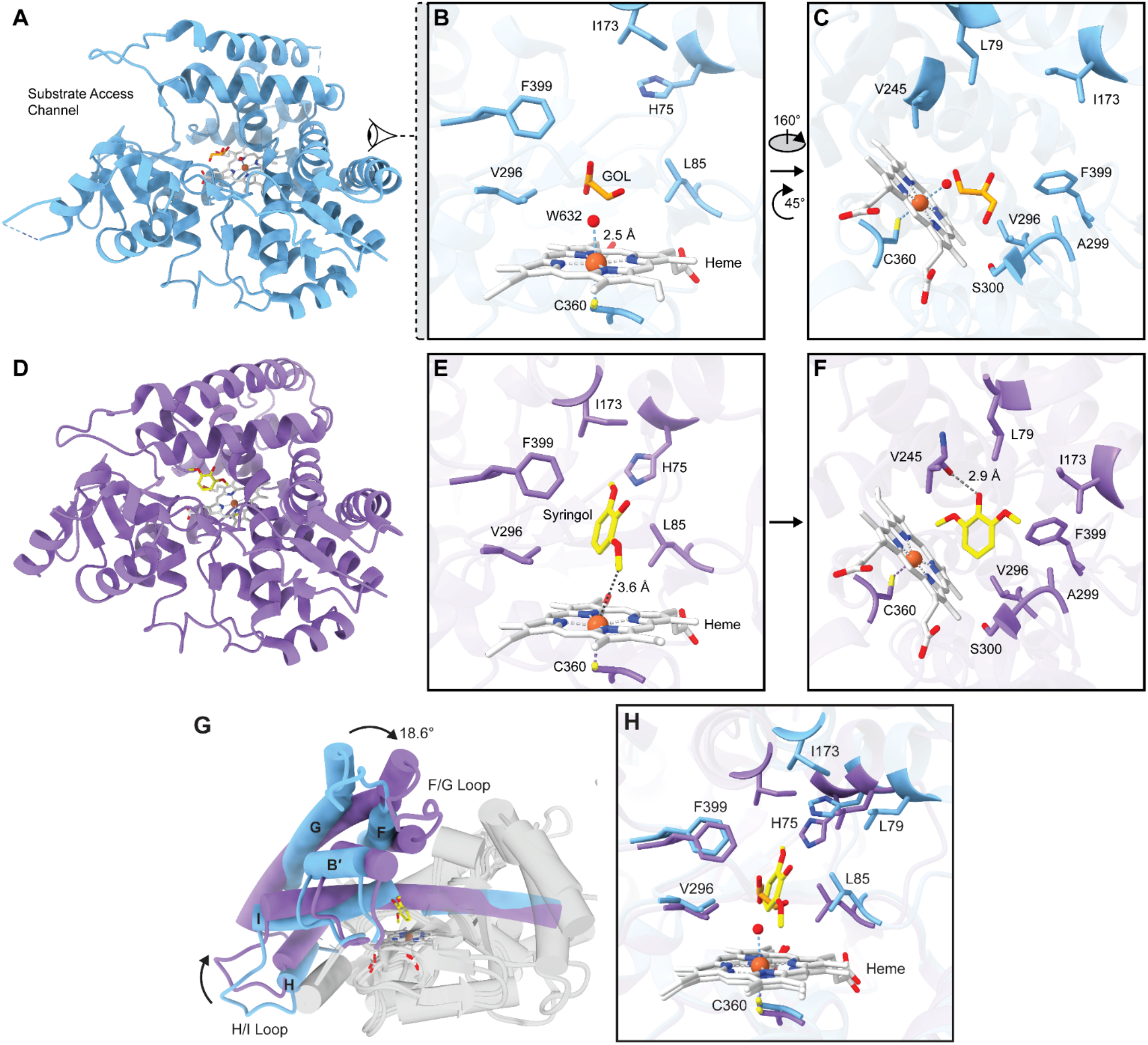
X-ray crystal structures of SyoA in the open substrate-free and closed substrate-bound states. Ribbon diagram showing the overall architecture of SyoA in the (A) open conformation (PDB: 8u09, blue) highlighting the positions of the heme (white), sixth axial water (red) and glycerol (GOL - orange/red). (B–C) Active site of the open conformation showing the key active site residues highlighting the solvent molecules and Fe-H_2_O distance (blue dashed line). Ribbon diagram showing the overall architecture of SyoA in the (D) closed conformation (PDB: 8u19, purple) showing the positions of the heme (white) and substrate syringol (yellow/red). (E–F) Active site of the closed conformation highlighting key active site residues; black dashed lines illustrate the distance of the 2-methoxy group of syringol from the Fe center, and the hydrogen bonding distance between the 1-hydroxyl group of syringol and the backbone carbonyl of Val245. The view of the active site in the right panel is made by rotating the left panel ∼160° clockwise about the *y*-axis, and ∼45° clockwise about the *z*-axis. Superposed structures of the open and closed form of SyoA showing the (G) overall fold and (H) active site. Only the helices that have an average Cα RMSD of 3 Å or greater are colored.

SyoA exhibits the characteristic single-domain trigonal prism shape, consisting of sixteen α-helices and four β-sheets, with the central heme iron coordinated by Cys360 (Fig. 2 A and D; SI Appendix, Fig. S1). The substrate-free enzyme adopts an open conformation, with a water-filled channel that exposes the active site and heme to solvent (Fig. 2 A, B and C; SI Appendix, Fig. S2A). The substrate access channel consists of a network of 13 hydrogen-bonded waters (SI Appendix, Fig. S3). On the distal side of the heme, an axial water is coordinated ∼2.5 Å from the heme iron (Fig. 2B). Furthermore, unidentified electron density was observed in the active site that was too large for water and too small for syringol. The best candidate for the densities above the heme and Phe399 was glycerol from the cryoprotectant, while density near His75 was consistent with acetate present in the crystallization conditions (SI Appendix, Fig. S3).

In contrast, the syringol-bound structure of SyoA adopts a closed conformation with the heme and substrate buried within a closed active site (Fig. 2D; SI Appendix, Fig. S2B). The active site of SyoA is predominantly hydrophobic, consisting of His75, Leu79, Leu85, Ile173, Val245, Val296, Ala299, Ser300, and Phe399 (Fig. 2 E and F). Syringol is situated in the active site, with one methoxy carbon positioned towards the heme ∼3.6 Å from the iron, and the other methoxy group positioned adjacent to His75 and Ile173 (Fig. 2E). Moreover, the aromatic ring of syringol is held between Leu85 and Phe399 with the hydroxyl group of syringol hydrogen-bonded to the carbonyl of Val245 (Fig. 2 E and F). The network of water molecules observed in the substrate-free structure are not present in the substrate-bound form.

The conformational change between the open and closed state of SyoA is characterized by an ∼19° displacement of helices B′, F, G and H; and the F/G and H/I loops, via a hinge-like mechanism (Fig. 2G; SI Appendix, Fig. S4A and Table S2). Displacements are observed for residues His75 (2.5 Å) and Leu79 (2.6 Å) of the B′-helix and Ile173 (4.4 Å) of the F-helix, which directly interact with one of the methoxy moieties of syringol (Fig. 2H; SI Appendix, Table S3). The G/H-loop, comprising residues 206–221, was not modelled in the open conformation due to poor resolution of the electron density. The average *B*-factors were highest surrounding the G/H-loop and were significantly lower in the closed state, consistent with stabilization of this region upon substrate binding (SI Appendix, Fig. S4B). Clear electron density was observed in the closed conformation enabling building and refinement of the G/H-loop in the bound state.

The closest structural homologue to SyoA is GcoA (PDB: 5ncb) (2); the aligned structures reveal that relative to GcoA, there is additional space in the active site of SyoA, consistent with SyoA being permissive of binding to S-lignin aromatics which contain an additional methoxy group compared to G-lignin aromatics (SI Appendix, Fig. S5 and S6). A triad of phenylalanine residues (Phe75, Phe169, and Phe395) is critical in positioning the aromatic ring of guaiacol in the active site of GcoA (2). SyoA does not contain this triad of phenylalanine residues; instead, Leu79, Ile173 and Phe399 occupy equivalent positions in the active site (SI Appendix, Fig. S6 A and B). Unique to the active site of SyoA is His75 (the equivalent residue in GcoA is Gly71), which partially fills the void in the active site occupied by larger aromatic residues in GcoA (SI Appendix, Fig. S6A).

### Active site binding of para-substituted S-lignin aromatics

Substrate binding to P450s commonly induces a shift from a low to high-spin ferric state. The low-spin (LS) state is characterized by heme coordination of 6 ligands (Soret maximum at ∼419 nm), including the active site water; the transition to a high-spin (HS) state (Soret maximum at ∼390 nm) occurs when the substrate displaces this water. Previously, our analysis showed that SyoA preferentially binds syringol over guaiacol (4). The high-resolution crystal structure of syringol-bound SyoA reveals additional space in the active site at the substrate C4 carbon – *para* to the hydroxyl group of syringol (Fig. 3 A and B). Consequently, we investigated binding of *para*-substituted S-lignin aromatics to SyoA. Among the tested S-type aromatics, only two induced a significant type I shift in the UV-vis absorbance spectrum, with 4-methylsyringol inducing a ∼70% shift to the HS form; and 4-allylsyringol a ∼20% shift (SI Appendix, Fig. S7). Other S-type aromatics including syringic acid, acetosyringone, sinapic acid, and sinapyl alcohol did not induce a type I shift (SI Appendix, Fig. S7). Strong absorbance of syringaldehyde in the visible spectrum from 420–390 nm introduced ambiguity in the binding titrations of this substrate; therefore, the Q-bands at 568 and 535 nm were used to interrogate binding (SI Appendix, Fig. S8). Addition of syringaldehyde to SyoA causes a decrease in the Q-bands, and an increase in the charge transfer band at ∼640 nm indicative of ligand binding in the active site. Binding analysis showed that SyoA binds 4-methylsyringol with the highest affinity (*K*_D_ = 7.2 ± 0.2 μM), followed by syringol (*K*_D_ = 16 ± 1 μM), syringaldehyde (*K*_D_ = 200 ± 12 μM) and 4-allylsyringol (*K*_D_ = 206 ± 11 μM) (SI Appendix, Fig. S9).

**Fig. 3:**
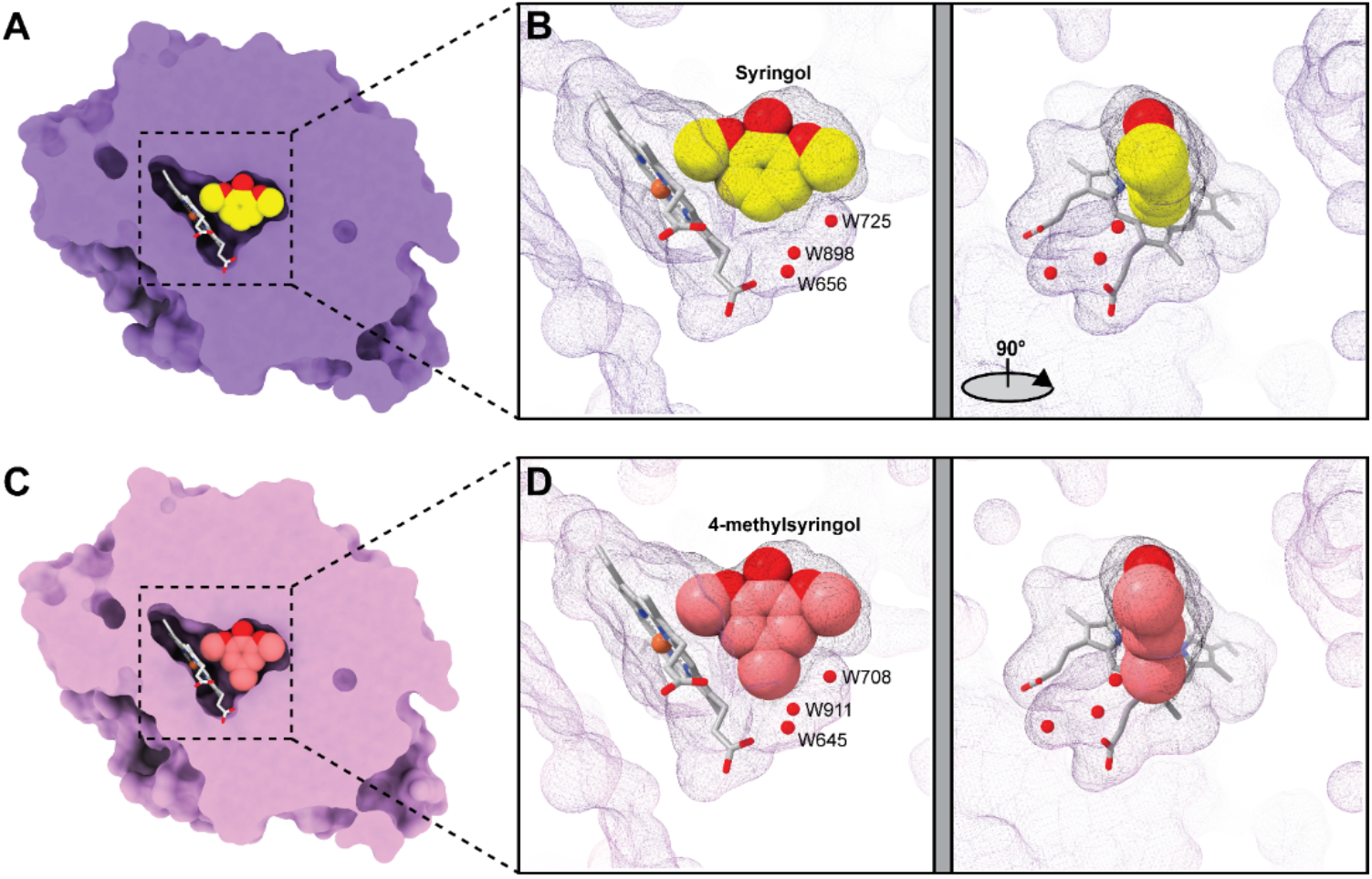
Comparison of active site volumes in the closed substrate-bound state. Surface rendering of (A) a slab view of the active site of SyoA (purple) bound to syringol (yellow/red) and (B) wireframe surface of the active site pocketshowing the position of the heme (white), syringol (yellow/red) and waters (small red spheres) that occupy the pocket. Surface rendering of (C) a slab view of the active site of SyoA (pink) bound to 4-methylsyringol (light red/red) and (D) wireframe surface of the active site pocket showing the heme (white), 4-methylsyringol (light red) and waters (small red spheres). Surface renders (A, C) are clipped through the center of SyoA at the position of the substrate. The second image (B, D) is rotated 90° clockwise relative to the first pane.

Crystallization of SyoA was subsequently conducted with the addition of 4-methylsyringol. The atomic-resolution crystal structure of SyoA bound to 4-methylsyringol (PDB: 8u1I) was solved at 1.12 Å; no significant differences were observed in the active site, nor in the position of the substrate, with an all-atom RMSD of 0.35 Å between the syringol and 4-methylsyringol-bound structures (Fig. 3 C and D; SI Appendix, Fig. S2C, S10, and Table S1). The 4-methyl group occupies the vacant space observed in the syringol-bound structure, with the remaining space occupied by three water molecules which are conserved in both structures (Fig. 3 C and D).

### SyoA catalyzes *O*-demethylation of S-lignin aromatics using the peroxide shunt

The CYP255 members GcoA and AgcA catalyze the *O*-demethylation of G-lignin aromatics, as a two-component system, using their respective three-domain redox partners (GcoB and AgcB). To determine the catalytic range of SyoA, we first investigated the *O*-demethylation of S-lignin aromatics using the monooxygenase pathway with the predicted redox partner SyoB, the encoding gene being adjacent that of SyoA (SI Appendix, Fig. S11). In contrast to GcoB and AgcB, SyoB consists of a predicted N-terminal FMN binding domain, NADH binding domain and a 2Fe-2S ferredoxin domain making it more structurally similar to the phthalate dioxygenase reductase family (30). Unlike the other CYP255 members, the reconstituted SyoAB monooxygenase system did not *O*-demethylate syringol, 4-methylsyringol, or syringaldehyde (SI Appendix, Fig. S12 and S13). Although NADH was consumed in the reactions with syringol, HPLC analysis identified that SyoB could react with syringol to form 2,6-dimethoxyhydroxyquinone in the absence of SyoA (SI Appendix, Fig. S12).

We next investigated the *O*-demethylation of syringol, 4-methylsyringol, syringaldehyde and 4-allylsyringol using the peroxide shunt pathway. Reactions with syringol, 4-methylsyringol and syringaldehyde yielded singly demethylated products and formaldehyde as a by-product (Fig. 4; SI Appendix, S14). We observed no evidence that SyoA could do two successive demethylation reactions to produce pyrogallols. The *O*-demethylation of syringol yielded 3-methoxycatechol (143 ± 12 μM, total turnover number (TTN) 143) as the major product, with ∼65% of the substrate consumed over 60 min (Fig. 4; SI Appendix, Fig. S15 and S19A). The demethylation of 4-methylsyringol occurred at the fastest rate, with 100% consumption of the substrate over 60 min (SI Appendix, Fig. S16A). However, none of the expected product was observed due to its overoxidation. To prevent product overoxidation, the substrate concentration was increased to 5 mM and the amount of H_2_O_2_ reduced to 4 mM. HPLC analysis revealed the formation of a major product at 10.1 min which was confirmed as 3-methoxy-5-methylbenzene-1,2-diol by GC-MS (299 ± 2 μM, TTN 299) (SI Appendix, Fig. 4, Fig. S16B and Fig. S19B). Syringaldehyde was demethylated to 5-hydroxyvanillin (308 ± 27 μM, TTN 308) (Fig. 4; SI Appendix, Fig. S17A). The reaction with syringaldehyde also liberated 2-hydroxy-6-methoxy-1,4-benzoquinone and 2,6-dimethoxy-1,4-benzoquinone due to oxidation of the aldehydes by hydrogen peroxide via a Dakin oxidation reaction (31) (SI Appendix, Fig. S17 B and C, S19C and Scheme S1). The oxidation of 4-allylsyringol by SyoA yielded several metabolites in low yield as observed by HPLC (SI Appendix, Fig S18).

**Fig. 4.**
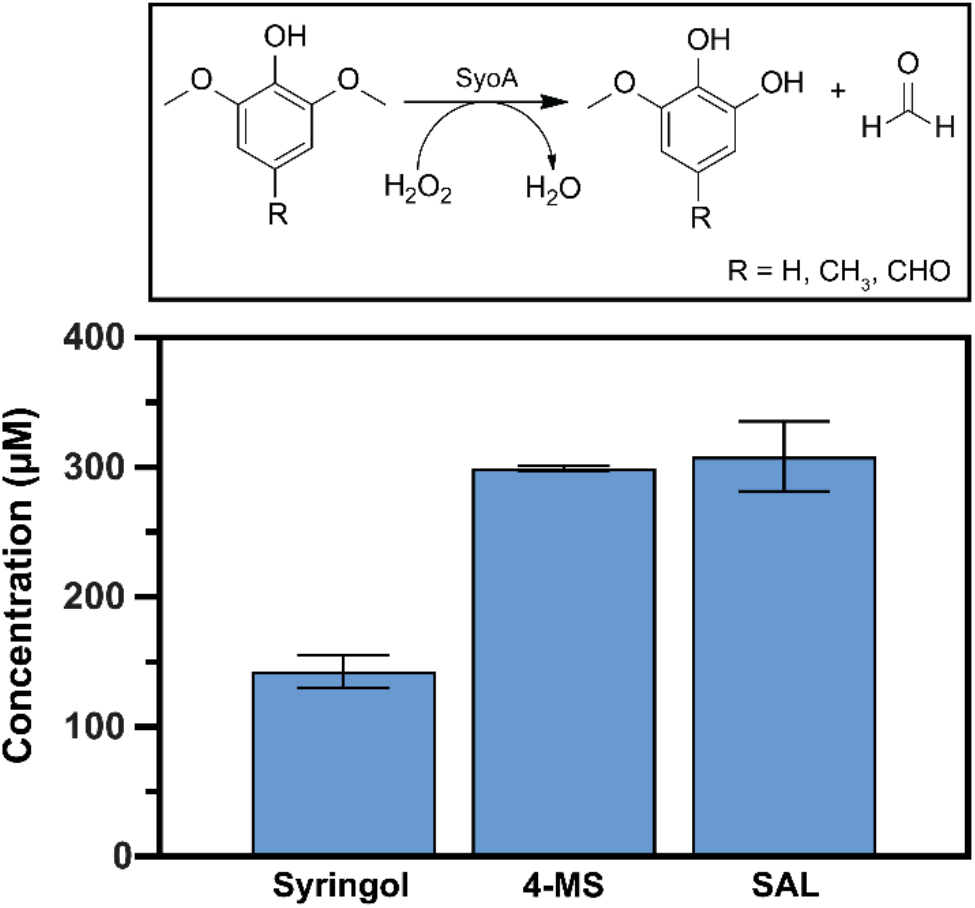
*O*-demethylation of S-lignin aromatics by SyoA using the peroxide shunt. Scheme showing the *O*-demethylation of S-lignin aromatics to form catechols and formaldehyde using the peroxygenase activity of SyoA. Concentration of the singly demethylated metabolites produced during the *O*-demethylation of syringol, 4-methylsyringol (4-MS) and syringaldehyde (SAL). Reactions were carried out using 1 µM SyoA in 50 mM Tris pH 7.5, 30°C, 60 min. The substrate concentration was 0.5 mM for syringol and SAL and 5 mM for 4-MS. Note the concentration of H_2_O_2_ was decreased to 4 mM for the 4-methylsyringol reaction, which had a TTN >500 for the equivalent 10 mM H_2_O_2_ reaction. Error bars indicate the SD from the mean of 3 replicates.

### Structural analysis and mutagenesis of the I-helix catalytic site residues

The I-helix of P450 enzymes contains residues important for oxygen activation and catalysis (32). We previously identified that the I-helix of the CYP255 family does not have the conserved acid-alcohol pair believed to be responsible for the monooxygenase function of most P450 enzymes (4). The crystal structures of SyoA and GcoA reveal that the glutamine-glutamate pair (Gln252 and Glu253 SyoA; Gln248 and Glu249 GcoA) are observed in a structurally equivalent position to the acid-alcohol residues of most P450 enzymes and are likely to be important for the peroxygenase activity of the CYP255 family (Fig. 5 A, B and C).

**Fig. 5.**
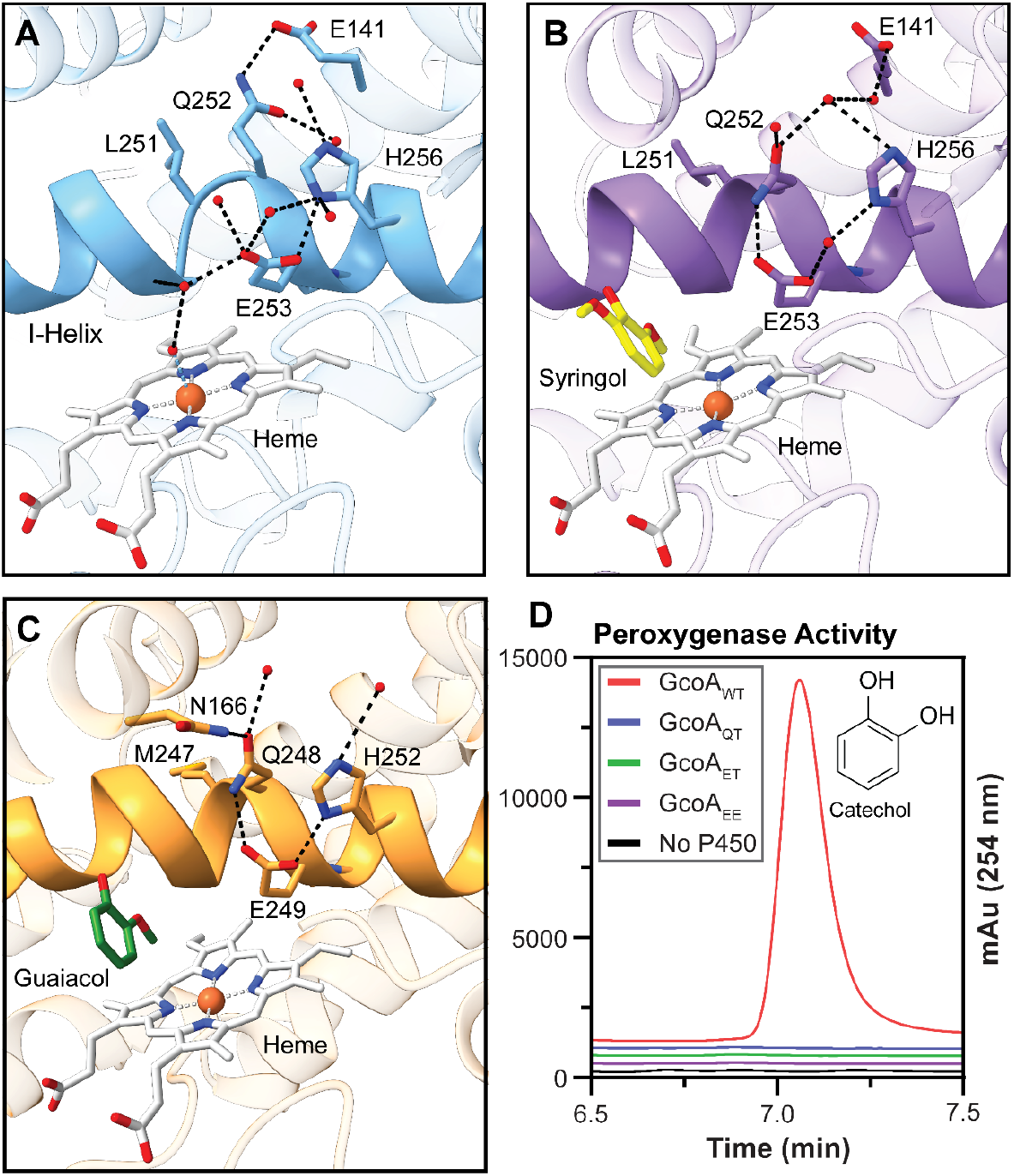
I-helix residues in the catalytic site. Ribbon and stick diagrams showing the I-helix of (A) SyoA substrate-free (PDB: 8u09), (B) SyoA syringol-bound (PDB: 8u19) and (C) GcoA guaiacol-bound (PDB: 5ncb). The I-helix is highlighted showing the key catalytic residues Gln252 and Glu253 and hydrogen bonding interactions (black dashed line) to surrounding residues and water molecules. (D) HPLC analysis of the hydrogen peroxide driven *O*-demethylation of guaiacol by mutants of GcoA including Q248E, E249T, and combinations thereof. Despite being folded, and competent for substrate binding, mutation of Gln248 and/or Glu249 resulted in complete loss of peroxygenase activity.

In the substrate-free state of SyoA, the I-helix exhibits a break in the α-helical structure from Gly249 to Gln252 due to the backbone amide of Glu253 forming a hydrogen bond with the carbonyl oxygen of Gly250 (Fig. 5A). In the syringol-bound complex, the break is resolved due to the backbone amide of Glu253 forming a hydrogen-bond with the carbonyl of Gly249 instead (Fig. 5B). This conformational change affects the position of the glutamine–glutamate pair. In the substrate-free structure Gln252 is hydrogen-bonded to Glu141 of the E-helix, while Glu253 forms several interactions with surrounding solvent molecules and His256 (Fig. 5A). Upon substrate binding, Gln252 forms a hydrogen bond to Glu253, moving closer to the heme (Fig. 5B). Additionally, the direct hydrogen bond between Glu253 and His256 is disrupted by a water molecule. The I-helix of GcoA (PDB: 5ncb) adopts a similar conformation to SyoA, however, Gln248 hydrogen bonds to Asn188 of the F-helix and Glu249 directly interacts with His252 (Fig. 5C). In the substrate-bound structures the glutamic acid is located ∼7.4 Å (SyoA) or ∼8.1 Å (GcoA) from the heme iron and ∼6.0 Å (SyoA) or ∼6.8 Å (GcoA) from the 2-methoxy carbon.

To investigate whether the non-canonical I-helix residues of the CYP255 family are important for peroxygenase activity, we performed site-directed mutagenesis on GcoA at Gln248 and Glu249. GcoA was selected as the impact on both the monooxygenase and peroxygenase activity could be assessed. We investigated the double mutant GcoA_ET_ which converted the QE residues (of GcoA_QE_) to those expected in a typical monooxygenase enzyme. In addition, we studied the importance of each individual residue on the activity using the GcoA_EE_ and GcoA_QT_ mutants. The ferric, ferrous and ferrous-CO spectra of each mutant were typical for P450 enzymes indicative of successful folding and heme incorporation (SI Appendix, Fig. S20A). Additionally, UV-Vis analysis demonstrated no loss of substrate binding to guaiacol (SI Appendix, Fig. S20B). Mutation of either Gln248 or Glu249 resulted in complete loss of peroxygenase activity in GcoA (Fig. 5D). Furthermore, a significant reduction in the monooxygenase activity was observed for GcoA_EE_ and GcoA_ET_ with complete loss in activity for the GcoA_QT_ mutant (SI Appendix, Fig. S21).

## Discussion

In contrast to chemical catalytic approaches, the *O*-demethylation of lignin aromatics using biocatalysis is of great interest as it offers an environmentally friendly method for converting lignin to high-value compounds (33, 34). S-lignin specific *O*-demethylases are rare, limiting the bioconversion of lignin from hardwoods and grasses. In this study, we determined the X-ray crystal structures of CYP255 SyoA in the open substrate-free and closed substrate-bound states and show that, *in vitro*, SyoA only catalyzes *O*-demethylation using the peroxide shunt pathway.

Compared to GcoA, SyoA has additional space in the active site allowing for binding of larger S-lignin aromatics. Specifically, SyoA contains an isoleucine (Ile173) at the equivalent position of a key phenylalanine residue (Phe169) present on the F-helix, which has been shown to prevent productive binding of syringol in GcoA due to a steric clash with the second methoxy group (11). Mutagenesis of this residue in GcoA to an alanine increases specificity for syringol (11) demonstrating that Ile173 in SyoA and Phe169 in GcoA may be responsible for determining specificity between S-lignin and G-lignin aromatics, respectively. Compared to GcoA, the active site of SyoA contains additional space at the *para* position allowing for binding of *para-*substituted aromatics (4-methylsyringol, syringaldehyde and 4-allylsyirngol), due to the presence of an active site serine (equivalent position in GcoA features a threonine). Syringaldehyde is a major product of oxidative lignin depolymerization (35, 36), while syringol and 4-methylsyringol are more commonly observed during lignin pyrolysis (36, 37), making SyoA a suitable catalyst for aromatics derived from different methods of lignin depolymerization.

Various P450 enzymes are known to undergo conformational changes from open to closed states upon substrate binding (38). In this work, we provide structural evidence that SyoA switches from an open to closed conformation in the presence of substrate, with relocation of the F, G and B′ helices to establish the closed hydrophobic active site of SyoA. The results described here support molecular dynamics (MD) simulations that show closing of the active site of GcoA in response to substrate binding (2). Contrary to these observations, MD simulations predict AgcA (CYP255A1) prefers a closed conformation with smaller substrates but adopts an open conformation with larger substrates (39).

Like most P450 monooxygenase enzymes, GcoA and AgcA catalyze substrate oxidation using their respective redox partners, NAD(P)H and O_2_. Interestingly, SyoA with its predicted redox partner SyoB demonstrated no *O*-demethylation of syringol, 4-methylsyringol and syringaldehyde. However, low levels of SyoB-dependent oxidation of syringol to 2,6-dimethoxyhydroxyquinone was observed. The presence of redox active quinones can result in NADH oxidation, transferring the electrons via the redox partners to the quinone, which can then react with O_2_ to ultimately form H_2_O_2_. This results in uncoupling of electron transfer from the P450 activity, explaining the oxidation of NADH without product formation (40). This phenomenon has also been observed for VanB and syringic acid (10) indicating that some lignin aromatics may lead to uncoupling via redox cycling. However, SyoA and GcoA can efficiently use the peroxide shunt pathway to catalyze *O*-demethylation of their substrates—an uncommon feature of P450s (4). Using the peroxygenase activity of SyoA, we were able to show *O*-demethylation of syringol, 4-methylsyringol and syringaldehyde. To our knowledge this is the first report of an enzyme that can demethylate S-lignin aromatics including 4-methylsyringol and syringaldehyde— which present as key bottlenecks to biological lignin valorization. Furthermore, atomic resolution structural insights also offer scope for engineering the active site of SyoA to accommodate other S-lignin aromatic substrates including syringic acid or 3-*O*-methylgallic acid. Additionally, the peroxygenase activity of the CYP255 family offers a cheap and clean method for demethylation compared to traditional monooxygenases, which require redox partners and expensive nicotinamide cofactors.

We postulate that the CYP255 family may have evolved hybrid monooxygenase/peroxygenase enzyme activity for several reasons. Exposing lignin to light facilitates production of hydrogen peroxide (41, 42). Interestingly, many enzymes involved in the depolymerization of lignin also use H_2_O_2_ as a cosubstrate, suggesting these peroxygenases may have evolved to use hydrogen peroxide as a ‘free’ catalytic driver, removing the reactive oxygen species and thereby protecting the P450 from oxidative stress (43). Additionally, as shown by the reaction of syringol with SyoB, CYP255 enzymes may have evolved peroxygenase activity to avoid uncoupling via redox cycling by quinones. Hydroquinones including 2-methoxyhydroquinone (MBQH_2_) and 2,6-dimethoxyhydroxyquinone (DBQH_2_) can be derived from products of lignin degradation (44, 45). Thus, the CYP255 family may have evolved hybrid monooxygenase/peroxygenase enzyme activity as a mechanism to minimize uncoupling in the presence of quinones, using the hydrogen peroxide generated to drive catalysis.

High-resolution structural analysis of the I-helix of SyoA has revealed the conserved acid-alcohol pair—typically found in P450 monooxygenases—is replaced with an amine-acid pair (glutamine and glutamate). Interestingly, peroxidases and peroxygenases contain amino acids capable of acid-base chemistry (histidine, aspartate, or glutamate) on the distal side of the heme (22, 29). We performed site-directed mutagenesis on the amine-acid pair of GcoA (Gln248 and Glu249) and demonstrated the importance of the non-canonical I-helix residues for both the monooxygenase and peroxygenase function. We hypothesized that mutating the amine-acid pair to the conserved acid-alcohol pair (GcoA_QT_ and GcoA_ET_) would improve monooxygenase activity and impair peroxygenase activity. However, all mutants lost the ability to use the peroxide shunt pathway with significant reduction in the monooxygenase activity also observed, demonstrating the amide-acid pair is important for both oxygen and hydrogen peroxide activation in the CYP255 family. These results demonstrate that the CYP255 family may oxidize substrates using an alternative mechanism compared to the majority of P450s with the conserved acid-alcohol pair. Deploying the I-helix of the CYP255 family may enable the retrofitting of peroxygenase activity into other P450s, making biocatalysis more favorable for larger scale applications. Indeed, the peroxygenase activity of CYP119 (46) and CYP199, CYP154C8 and CYP102 (47) was improved by mutating the acid-alcohol pair and surrounding residues to resemble the I-helix of the CYP255 enzymes.

## Materials and Methods

### General

Chemical reagents were purchased from Sigma-Aldrich. HPLC grade solvents were from Sigma-Aldrich, Ajax Finechem and Chem-Supply. Enzymes, biochemical and molecular biology reagents were supplied by Sigma-Aldrich, New England Biolabs and Thermo Fisher Scientific. Crystallography reagents, MicroMounts, MicroLoops and MicroTools were from Hampton Research and Molecular Dimensions and the NVH oil from (Cargille Laboratories). Substrate and product stocks were prepared in DMSO. IS (internal standard) stock solutions were prepared in EtOH.

### Gene cloning

A pET3a vector containing a codon optimised gene encoding SyoA was obtained using 3-fragment Gibson isothermal assembly. The pET26 vector containing GcoA was used as previously described (4). GcoA mutants (GcoA_EE_, GcoA_QT_, GcoA_ET_) were purchased (Twist Bioscience) cloned in the pET28a vector. SyoB and GcoB were cloned into the pET29b vector by PCR of gBlocks purchased from (IDT) and restriction digest using *Nde*I and *Hind*III. Details are provided in SI Appendix.

### Protein expression and purification

SyoA, GcoA and GcoA mutants (GcoA_EE_, GcoA_QT_ and GcoA_ET_) were expressed and purified using previously described methods (4). SyoB and GcoB were transformed into *E. coli* BL21 (DE3) cells and grown in LB media containing kanamycin (50 µg/ml) at 37 °C to an optical density OD_600_ of 0.6. Gene expression was induced by addition of 0.1 mM isopropyl β-D-1-thiogalactopyranoside (IPTG), followed by incubation at 20 °C and 80 rpm. Additionally, 1% v/v ethanol, 0.02% v/v benzyl alcohol, 1 mM L-cysteine and 0.5 mM ferric ammonium citrate were added prior to induction. Cells were harvested by centrifugation, resuspended in (50 mM Tris, pH 7.5, 500 mM NaCl, 20 mM imidazole, 1 mM DTT) for SyoB and (50 mM Tris, pH 7.5, 50 mM NaCl, 1 mM DTT) for GcoB, and lysed by sonication. SyoB was purified by immobilized metal-affinity chromatography (IMAC) and GcoA by ion-exchange chromatography. Details provided in SI Appendix.

### Protein crystallization and structure determination

For crystallization, the N-terminal His_6_ tag was cleaved using His_6_-TEV protease and isolated from His_6_-tagged SyoA and TEV protease by IMAC. SyoA was further purified by size exclusion chromatography (HiLoad 16/600 Superdex 200 pg, Cytiva) and concentrated to approximately 15 mg mL^-1^ in 20 mM Tris pH 7.5 by ultrafiltration. Crystals were grown by sitting drop vapor diffusion at 289 K using 1 µL of protein with 1 µL of reservoir solution, including PEG/Ion I&II, Index, and PGA screens. SyoA in the substrate-free state crystallized within 1 week in conditions comprising 0.25 mM syringol (from a 50 mM stock in DMSO), 0.1 M sodium acetate pH 5, 5% w/v γ-PGA (γ-polyglutamic acid) and 12% w/v PEG 8000. Crystals were cryoprotected in reservoir solution supplemented with 25% v/v glycerol prior to flash cooling in liquid nitrogen. SyoA in the substrate-bound state crystallized within 1 week in conditions comprising 20 mM substrate (from a 200 mM stock in 40% DMSO), 0.2 M magnesium nitrate hexahydrate pH 5.9, 20% w/v PEG 3350 or 0.2 M lithium nitrate pH 7.1, 20% w/v PEG 3350. Crystals were cryoprotected in NVH oil prior to flash cooling in liquid nitrogen.

Diffraction data were collected for the substrate-free and syringol-bound crystals of SyoA at the MX1 beamline at the Australian Synchrotron (48). Diffraction data were collected for the 4-methylsyringol-bound crystal of SyoA at the MX2 beamline at the Australian Synchrotron (49) 360° of data were collected using 0.1° oscillations at a wavelength of 0.9357 Å at 100 K. The data were autoprocessed, indexed and integrated using xdsme (50) in space group P2_1_2_1_2 for the substrate-free structure and P2_1_ for the substrate-bound structures. Data were scaled and merged using Aimless (51) and truncated to 1.98 Å, 1.26 Å and 1.12 Å for the substrate-free, syringol-bound and 4-methylsyringol-bound structures, respectively. Molecular replacement was performed using Phaser-MR (52) with an Alphafold2 (53) model of SyoA made using ColabFold (54) as the initial search model for the substrate-free structure, which was subsequently used to phase the syringol-bound model. The refined syringol-bound structure was then used as the search model to phase the 4-methylsyringol-bound data. The models were refined with Phenix.refine and built and improved in Coot (55) over multiple iterations. Data collection and refinement statistics are summarized in Table S1. Molecular graphics, structural alignments and RMSD calculations were performed with UCSF ChimeraX (56).

### Substrate binding analysis and dissociation constants

Substrate binding and the dissociation constant (*K*_*D*_) for SyoA and syringol, 4-methylsyringol, semialdehyde and 4-allyl syringol was determined as previously described (4) and detailed in SI Appendix.

### CO binding assays

CO binding assays for the GcoA mutants were performed as described previously (4) and described in detail in SI Appendix.

### *In vitro* NADH/O_2_-driven reactions

*In vitro* NADH/O_2_-driven reactions were performed at 30 °C in a total volume of 600–1200 μL. Reactions were performed in a quartz cuvette (with a 1 cm path length) and contained 1 μM SyoA or GcoA, 1 μM SyoB or GcoB, 100 μg mL^−1^ catalase (from bovine liver), 1 mM substrate and 320 μM NADH in oxygenated Tris buffer (50 mM, pH 7.5). The rate of NADH consumption was measured spectroscopically at 340 nm. Further details are provided in SI Appendix.

### *In vitro* H_2_O_2_-driven turnovers

*In vitro* H_2_O_2_ turnovers to assess the *O*-demethylation of syringol, 4-methysyringol and syringaldehyde by SyoA and guaiacol by GcoA, GcoA_EE_, GcoA_QT_ and GcoA_ET_ were performed as previously described (4) and detailed in SI Appendix.

### Formaldehyde Assay

Formaldehyde production was measured using a colorimetric Purpald assay (57). To determine the concentration of formaldehyde, 120 μL of the *in vitro* turnover mixture was mixed with 48 μL of 32 mM Purpald (dissolved in 2 M NaOH) and 432 μL 50 mM Tris, pH 7.5. The reaction was allowed to develop at room temperature by shaking for 60 min. The absorbance at 550 nm was recorded and the concentration of formaldehyde calculated using a calibration curve made with 0–400 μM formaldehyde. Reactions excluding the Purpald reagent were used as a baseline and the concentration of formaldehyde normalized against controls reactions without P450.

### Product analysis by HPLC

HPLC was performed as previously described (4) to identify the products formed during the H_2_O_2_-driven or NADH-driven reactions. Details provided in SI Appendix.

### Product analysis by GC-MS

*In vitro* H_2_O_2_ turnovers were performed as described above in 600 µL and analytes extracted once with 300 µL ethyl acetate. Samples were thoroughly mixed for 30 s by vortex. After phase separation, 180 µL of the organic layer was transferred to vials and analyzed by gas chromatography-mass spectrometry (GC-MS). Details provided in SI Appendix.

### Data Deposition

The atomic coordinates and structure factors have been deposited in the Protein Data Bank, https://www.PDB.org with accession codes 8u09 SyoA; 8u19 syringol-bound SyoA, and 8u1i 4-methylsyringol-bound SyoA.

## Author Contributions

A.C.H, T.D, K.E.S, S.G.B and F.W designed research; A.C.H, T.D, and S.G.B performed research; A.C.H, T.D, S.G.B and F.W analyzed data; A.C.H, K.E.S, S.G.B and F.W wrote the paper.

## Funding and Acknowledgments

This work was funded, in part, through Australian Research Council grants DP200102411 (to S.G.B and others) and DP230103062 (to F.W. and K.E.S.). This research was undertaken on the MX1 and MX2 beamlines at the Australian Synchrotron, part of Australian Nuclear Science and Technology Organization, and made use of the ACRF detector on MX2. We acknowledge financial support from the Australian Synchrotron to use this facility. A.C.H. was supported by Wine Australia, with levies from Australia’s grapegrowers and winemakers and matching funds from the Australian Government.

A.C.H and T.D were supported by Research Training Program (RTP) scholarships from the Australian government. F.W. was supported by the Ramsay Fellowship in Applied Science.

