## Supplementary Information for "The S-lignin *O*-demethylase SyoA: Structural insights into a new class of heme peroxygenase enzymes"

### Supplementary Materials and Methods

#### Gene cloning

A pET3a vector containing a codon optimized gene encoding SyoA was obtained using Gibson isothermal assembly (1), by 3-fragment assembly with a single polymerase chain reaction (PCR)-generated insert fragment containing the *syoA* gene and two plasmid backbone fragments. The *syoA* gene was PCR-generated from the previously described pET29 vector (2) using primers 3558 and 3559 (Table S6). The pET3a backbone fragments were PCR-generated using primers pairs 2281 and 3312 and 3497 and 1330. Primers 215 and 315 (T7 promoter and T7 terminator, respectively) were used to sequence the insert fragment. The pET26 vector containing GcoA was used as previously described (2).

GcoA mutants ($GcoA_{EE}$, $GcoA_{QT}$, $GcoA_{ET}$) were purchased (Twist Bioscience) cloned in the pET28a plasmid with an N-terminal 6 x His tag and Tobacco etch virus (TEV) cleavage site followed by a *Nde*I restriction site and the sequence for the gene of interest.

Codon optimized genes encoding SyoB (based on the NCBI reference sequence: WP_027930991) and GcoB were purchased as gBlocks from IDT. An *Nde*I restriction site was incorporated at the 5' end, followed by a double stop codon and *Kpn*I and *Hind*III restriction sites. PCR was performed on the gBlock DNA using primers RS0114865 5' and RS0114865 His 3' for SyoB and RS0109265 5' and RS0109265 His 3' for GcoB to add a 6 x His tag to the C-terminus of the gene and inserted into a pET29b vector using the *Nde*I and *Hind*III restriction sites. The final sequence was confirmed by DNA sequencing (Australian Genome Research Facility, Adelaide, Australia).

#### Protein expression and purification of SyoA, GcoA and GcoA mutants ($GcoA_{EE}$, $GcoA_{QT}$ and $GcoA_{ET}$)

The pET3a vector containing the *syoA* gene, the pET26 vector containing the *gcoA* gene and the pET28a vectors containing $gcoA_{EE}$, $gcoA_{QT}$ *or* $gcoA_{ET}$ were transformed into *Escherichia coli* BL21 (DE3) competent cells. The cells were plated on lysogeny broth (LB) agar containing ampicillin (100 µg/ml) for the pET3a vector and kanamycin (50 µg/ml) for the pET26 and pET28a vectors and incubated at 37 °C overnight (~16–20 h). Single colonies were used to inoculate a 50 mL LB starter culture containing the appropriate antibiotic and grown overnight (~16–20 h) at 37 °C with shaking (180 rpm). 2.5 L flasks containing 1 L LB with antibiotics were inoculated with 1:100 v/v starter culture and incubated, 37 °C with shaking (150 rpm). At an $OD_{600}$ of 2, 1% v/v ethanol and 0.02% v/v benzyl alcohol were added, and the cultures incubated at 18 °C with shaking 100 rpm. Additionally, 3 mL/L trace elements (Per liter: $Na_2EDTA$ (20.1 g), $FeCl_3.6H_2O$ (16.7 g), $CaCl_2.H_2O$ (0.74 g), $CoCl_2.6H_2O$ (0.25 g), $ZnSO_4.7H_2O$ (0.18 g), $MnSO_4.4H_2O$ (0.132 g), $CuSO_4.5H_2O$ (0.10 g)) were added for cofactor incorporation. After 30 min, protein expression was induced with addition of IPTG (isopropyl β-D-1-thiogalactopyranoside) to a final concentration of 0.1 mM, and the culture incubated overnight ~20 hours at 18 °C with shaking at 100 rpm. Cells were harvested by centrifugation (6200 x*g*, 15 min, 4 °C).

The cell pellet was resuspended in ~100 mL ice-cold lysis buffer (50 mM Tris, pH 7.5, 50 mM NaCl, 20 mM imidazole). Cells were lysed by sonication (30 cycles of 15 s on, 45 s off, 70% amplitude, 19 mm probe, Sonics Vibra-Cell) and cell lysate clarified by centrifugation (40,000 x*g*, 30 min, 4 °C). Proteins were purified by immobilized nickel affinity chromatography (HisTrap HP, 5 mL column Cytiva) using a linear gradient of 20–250 mM imidazole at a flow rate of 5 mL/min. The samples were buffer exchanged into 50 mM Tris, pH 7.5 by centrifugal ultrafiltration using a Vivaspin 20 centrifugal concentrator (Sartorius) with a 10 kDa molecular weight cut-off. Samples were further purified by anion exchange (HiTrap Q FF, 5 mL Cytiva) using a linear gradient of 0–500 mM NaCl at a flow rate of 5 mL/min. Purified protein was concentrated by ultrafiltration to <10 mL and stored in 40–50% v/v glycerol at -20 °C. Glycerol was removed before use using a PD-10 desalting column (Cytiva) equilibrated in 50 mM Tris, pH 7.5. The concentration of each protein was estimated using the following extinction coefficients: SyoA ($\varepsilon_{417nm}$ = 123 $mM^{-1}$ $cm^{-1}$)(2), GcoA ($\varepsilon_{417nm}$ = 119 $mM^{-1}$ $cm^{-1}$)(2).

**Protein expression and purification of SyoB and GcoB**

The pET29b vectors containing *syoB* or *gcoB* were transformed into *Escherichia coli* BL21 (DE3) competent cells. The cells were plated on lysogeny broth (LB) agar containing kanamycin (50 µg/ml) and incubated at 37 °C overnight (~16–20 h). Cells were grown as described above in 2.5 L flasks containing 1 L LB with antibiotics. At an $OD_{600}$ of 0.6, 1% v/v ethanol and 0.02% v/v benzyl alcohol were added, and the cultures incubated at 18 °C with shaking 80 rpm. Additionally, 1 mM L-cysteine and 0.5 mM ferric ammonium citrate were added for cofactor incorporation. After 30 min, protein expression was induced with addition of 0.1 mM IPTG (isopropyl β-D-1-thiogalactopyranoside), and the culture incubated overnight ~20 hours at 20 °C with shaking at 80 rpm. Cells were harvested by centrifugation (6200 x*g*, 15 min, 4 °C).

The cell pellet was resuspended in 100 mL ice-cold lysis buffer (50 mM Tris, pH 7.5, 500 mM NaCl, 20 mM imidazole, 1 mM DTT) for SyoB and (50 mM Tris, pH 7.5, 50 mM NaCl, 1 mM DTT) for GcoB. Cells were lysed by sonication (30 cycles of 15 s on, 45 s off, 70% amplitude, 19 mm probe, Sonics Vibra-Cell) and cell lysate clarified by centrifugation (40,000 x*g*, 30 min, 4 °C). SyoB was purified by immobilized nickel affinity chromatography (HisTrap HP, 5 mL column Cytiva) using a linear gradient of 20–250 mM imidazole at a flow rate of 5 mL/min. GcoB was purified by ion-exchange chromatography, using a DEAE Sepharose column. A linear gradient of 50–500 mM NaCl was used to elute the protein at a flow rate of 5 mL/min. The samples were buffer exchanged into 50 mM Tris, pH 7.5 by centrifugal ultrafiltration using a Vivaspin 20 centrifugal concentrator (Sartorius) with a 10 kDa molecular weight cut-off. Purified protein was concentrated by ultrafiltration to <10 mL and stored in 40–50% v/v glycerol at -20 °C. Glycerol was removed before use using a PD-10 desalting column (Cytiva) equilibrated in 50 mM Tris, pH 7.5. The concentration of SyoB and GcoB were estimated using the extinction coefficient ($\varepsilon_{423nm}$ = 25.2 $mM^{-1}$ $cm^{-1}$) (3).

**Substrate binding analysis and dissociation constants**

To determine substrate binding, SyoA or GcoA was diluted to ~3 µM in 50 mM Tris pH 7.5 in 600 µL and substrate added in 1 µL aliquots from a 100 mM stock in DMSO. The absorbance was measured between 600–250 nm on a Cary 60 UV-Vis Spectrophotometer (Agilent Technologies) until no further shift was observed**.**

To determine substrate binding affinity ($K_D$), SyoA was diluted to ~2–8 µM in 50 mM potassium phosphate buffer pH 7.4 in a volume of 2.5 mL. Aliquots (0.5–5 µL) of 1, 10, or 100 mM substrate stock solutions in DMSO were added using a 5 µL Hamilton syringe. The difference spectrum was recorded from 300–600 nm. Aliquots of substrate were added until no further spectral shift occurred and no more than 10 µL of each stock solution was added to avoid diluting the enzyme. Titrations were performed in triplicate. The peak-to-trough absorbance difference, ΔA ($A_{peak}$ – $A_{trough}$), was then plotted against substrate concentration. The ΔA was calculated using $A_{390-419\ nm}$ for all substrates except syringaldehyde ($A_{419\ nm}$). To obtain the dissociation constant the data were fitted to the hyperbolic (Michaelis-Menten) equation (Equation 1), where $K_D$ denotes the binding constant, [S] the substrate concentration, ΔA the peak-to-trough ratio, and $\Delta A_{max}$ the maximum peak-to-trough absorbance.

$$\Delta A = \frac{\Delta A_{max} \times [S]}{K_D + [S]} \qquad (1)$$

**CO binding assays**

The UV-Vis spectrum of ~3 µM GcoA, $GcoA_{EE}$, $GcoA_{QT}$ or $GcoA_{ET}$ (500 µL) in 50 mM Tris, pH 7.5 was recorded from 250–700 nm using a Varian Cary 300 UV-Vis Bio Spectrophotometer. A quartz cuvette with a 1 cm path length was used. A few grains of sodium dithionite were added and the spectrum of the reduced P450 was recorded. To form the P450-CO complex, CO was slowly bubbled through the solution for ~10s and the spectrum was recorded.

***In vitro* NADH/$O_2$-driven reactions**

*In vitro* NADH/$O_2$-driven reactions were performed at 30 °C in a total volume of 600–1200 µL. Reactions were performed in a quartz cuvette (with a 1 cm path length) and contained 1 µM SyoA or 0.5 µM GcoA, 1 µM SyoB or GcoB, and 100 µg $mL^{-1}$ catalase (from bovine liver) in oxygenated Tris buffer (50 mM, pH 7.5). This mixture was used to blank the spectrophotometer. NADH was added to a concentration of ~320 µM and the background rate of NADH air oxidation was measured spectrophotometrically at 340 nm (ε340 nm = 6.22 $mM^{-1}$ $cm^{-1}$). The reaction was initiated by addition of 1 mM substrate (from a 100 mM stock solution in DMSO) and NADH oxidation was monitored at 340 nm for the first ~10 min, and the reactions were then left at room temperature for ~1 h to ensure that the NADH was completely consumed. Control reactions were also performed in which the P450 was omitted from the turnover mixture.

***In vitro* $H_2O_2$-driven reactions**

*In vitro* $H_2O_2$ turnovers were performed at 30 °C and contained 1 µM P450, 0.5–5 mM substrate from a 100 mM stock dissolved in DMSO and 4–10 mM $H_2O_2$ for the SyoA reactions and 20 mM $H_2O_2$ for the GcoA reactions in Tris buffer (50 mM, pH 7.5) in a total volume of 600 µL. Each reaction was performed in triplicate. Reactions were incubated for 2 mins at 30 °C prior to addition of $H_2O_2$ to initiate the reaction ($H_2O_2$ was added from a 200 mM stock freshly prepared before each experiment from a 30% w/v stock). All reactions were quenched at 60 min with 10 µL of 10 mg $mL^{-1}$ catalase. Control reactions without P450 were performed to assess stability of substrates and products with $H_2O_2$.

**Product analysis by HPLC**

Analytical High Performance Liquid Chromatography (HPLC) was performed on a Shimadzu LC-20AD equipped with a Phenomenex Kinetex 5u XB-C18 100 Å column (250 mm x 4.6 mm, 5 µM), SIL-20A autosampler, CTO-20A, SPD-20A UV detector and CBM-20Alite communications module. Each sample and standard were injected at a volume of 20 μL. A gradient of 20–95% MeCN in water (with 0.1% TFA) over 30 minutes was used to elute the samples at a rate of 1 mL $min^{-1}$ and the eluate was monitored at 254 nm.

To prepare turnover mixtures for HPLC, 132 μL of the *in vitro* turnover mixture was mixed with 66 μL of MeCN and 2 μL of internal standard (IS, 10 mM 9-hydroxyfluorene in EtOH) and centrifuged at 15,800 x*g* for 3 mins to remove particulate matter. Calibration curves were constructed to quantify the product where available. Solutions of products with concentrations of 20, 50, 100, 200 and 500 μM were prepared for HPLC analysis in the same way as the turnovers. A plot of product peak area/IS peak area versus product concentration was then constructed.

**Product analysis by GC-MS**

*In vitro* $H_2O_2$ turnovers were performed as described above in 600 μL and analytes extracted once with 300 μL ethyl acetate. Samples were thoroughly mixed for 30 s by vortex. After phase separation, 180 μL of the organic layer was transferred to vials and analyzed by gas chromatography-mass spectrometry (GC-MS) using a Shimadzu GC-2010 equipped with a QP2010S GC-MS detector. 1 μL of sample was injected for analysis using a splitting ratio of 20. Separation of the analytes was done using a DB-5MS UI column (30 mm × 0.25 mm × 0.25 μm) at a flowrate of 1.4 mL $min^{-1}$ and the following temperature settings: Initial temperature of 100 °C held for 2 min, increased to 270 °C at 10 °C $min^{-1}$ and held at 270 °C for 6 min. The injection inlet temperature was 230 °C, ion source temperature 250 °C.

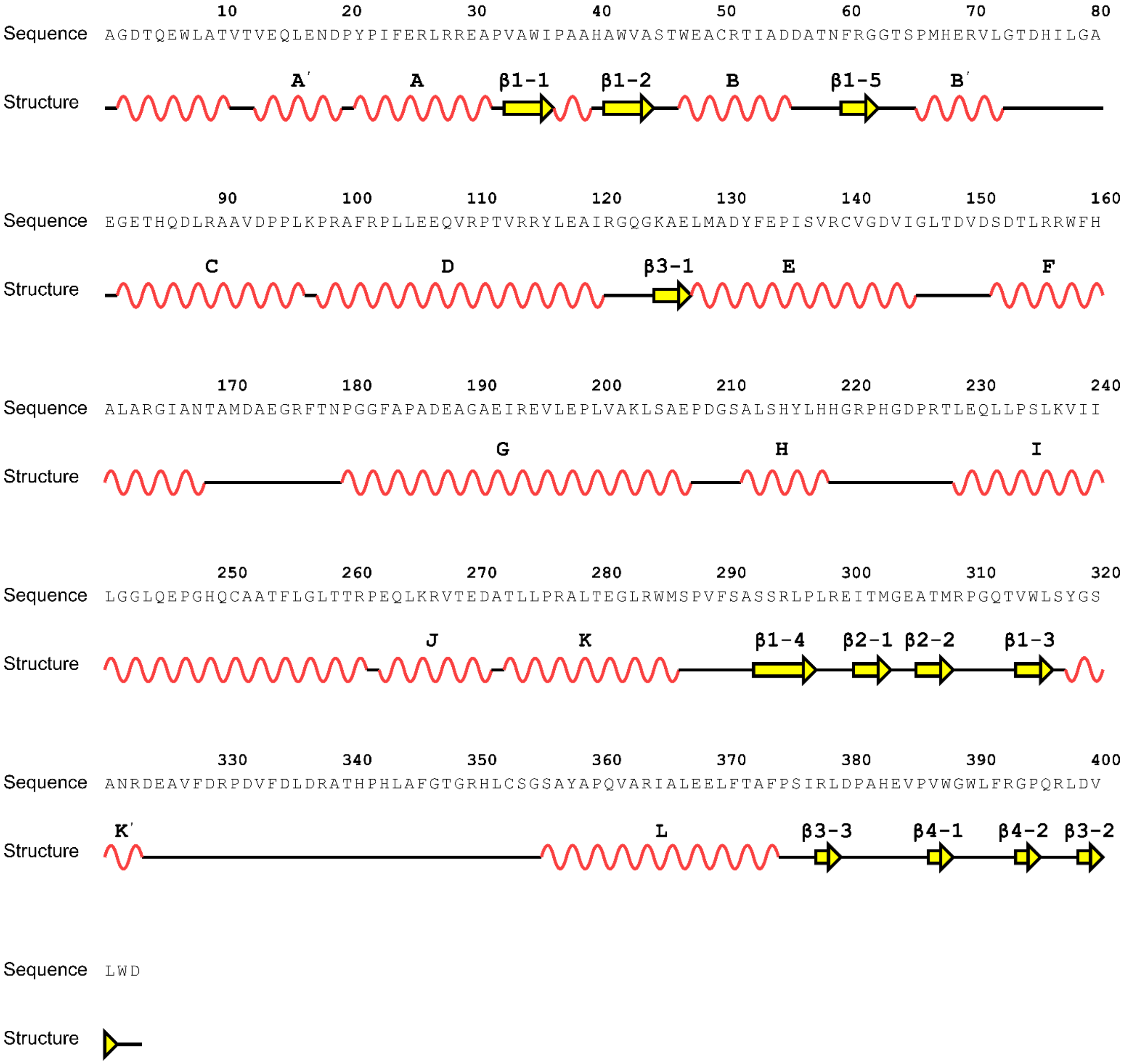


**Fig. S1.** Sequence and secondary structural topology of SyoA. The secondary structure of SyoA is illustrated below the sequence (α-helices, red wave; β-strands, yellow arrows). Visualization was created using the 2dSS webserver (4). Assignment of helices and β-sheets was done using SecStrAnnotator (5)

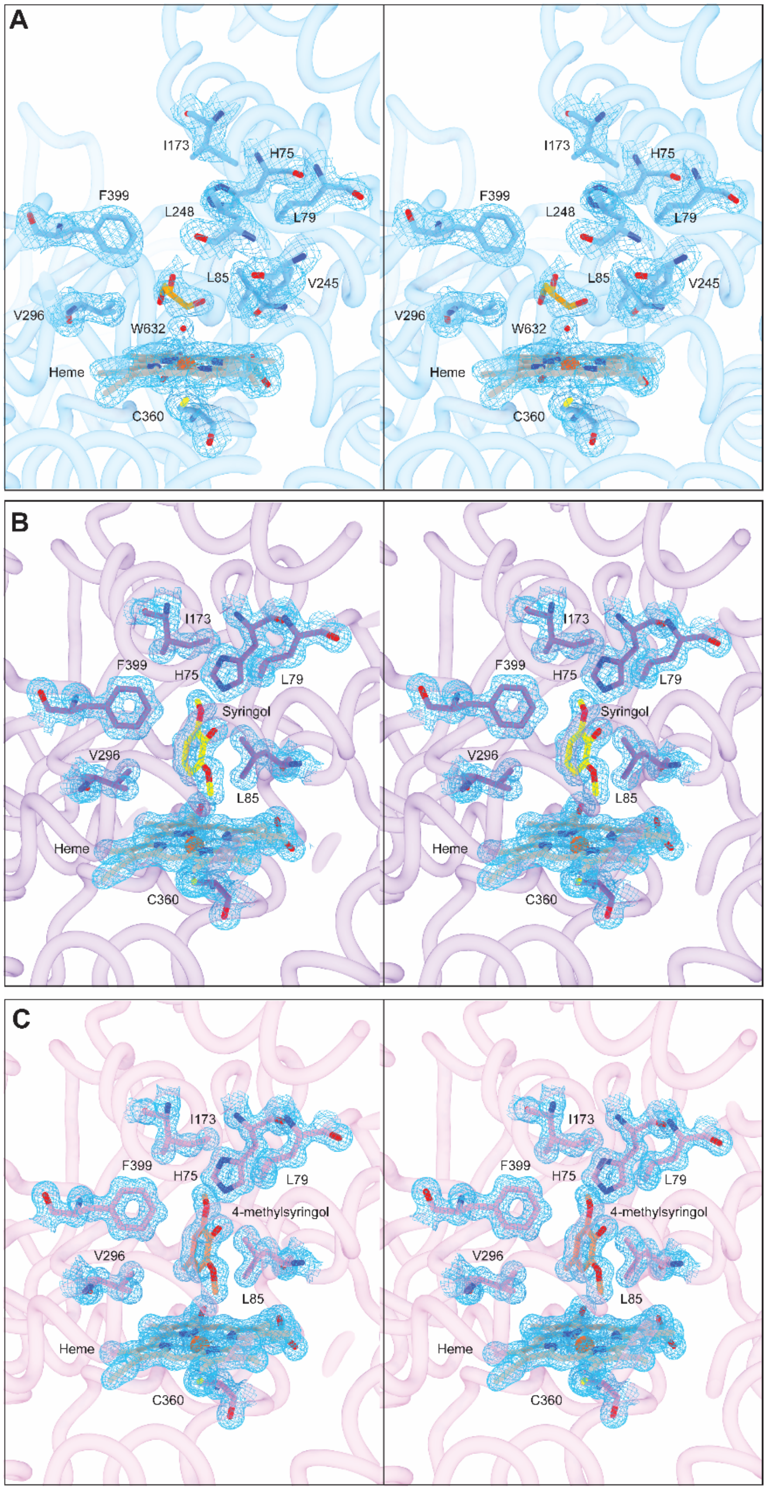


**Fig. S2.** Electron density of the active site of SyoA. Stereo view of the (A) substrate-free (PDB: 8u09) (B) syringol-bound (PDB: 8u19) and (C) 4-methylsyringol bound (PDB: 8u1i) structures of SyoA showing the electron density rendered around the substrate, heme group and the active site residues. Composite omit maps computed by Phenix refine using the simple method are shown in blue mesh contoured at 1σ (0.73, 0.81 and 0.84 e/Å$^{-3}$ for the substrate-free, syringol-bound and 4-methylsyringol-bound structures, respectively).

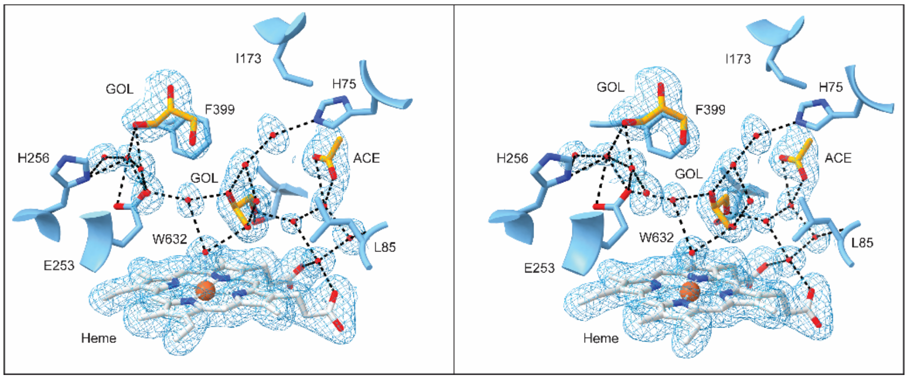


**Fig. S3.** Active site solvent channel of SyoA in the open conformation. Stereo view of the substrate-free structure of SyoA (PDB: 8u09) showing the electron density around the hydrogen-bonded solvent molecules (GOL, glycerol; ACE, acetate). Composite omit map contoured at 1σ (0.44 e/Å−3). The hydrogen bonds of the water network are shown as black dashed lines.

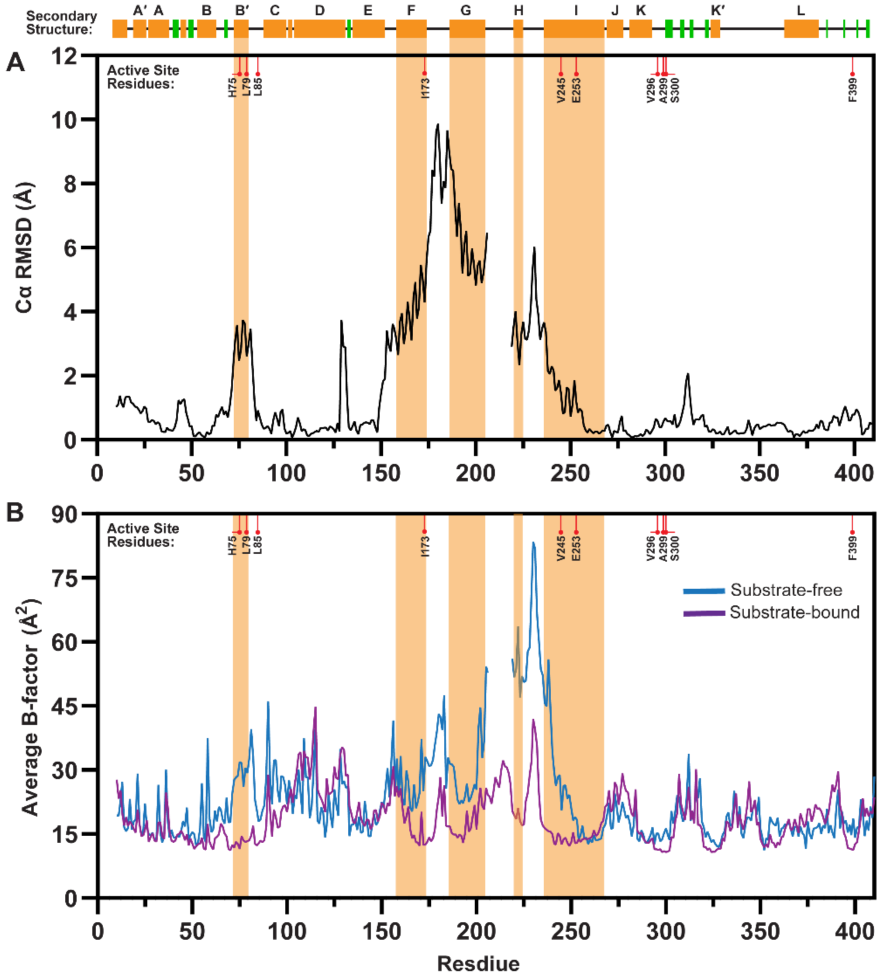


**Fig. S4.** Per residue (A) Cα RMSD (open vs closed) and (B) average B-factor for the substrate-free open (PDB: 8u09) and syringol-bound closed structures (PDB: 8u19). The secondary structure is denoted schematically above the graphs, with α-helices shown in orange and β-sheets in green. The B′, F, G, H and I-helix are highlighted in orange. Active site residues are denoted along the top of the *x*-axis. Residues 206–221 were unmodeled in the open structure, thus the Cα RMSD and average B-factor for the substrate-free structure could not be calculated.

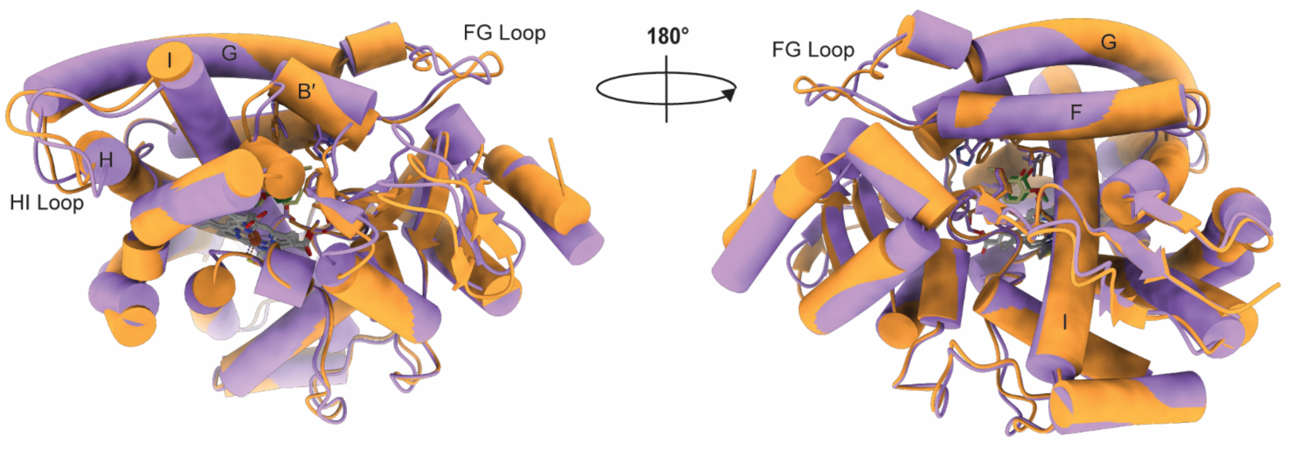


**Fig. S5.** Superposed structures of SyoA bound to syringol (PDB: 8u19, purple) and GcoA bound to guaiacol (PDB: 5ncb, orange) shown as cylinders and stubs (rendered in ChimeraX). The heme (white), syringol (yellow) and guaiacol (green) are shown as stick diagrams.

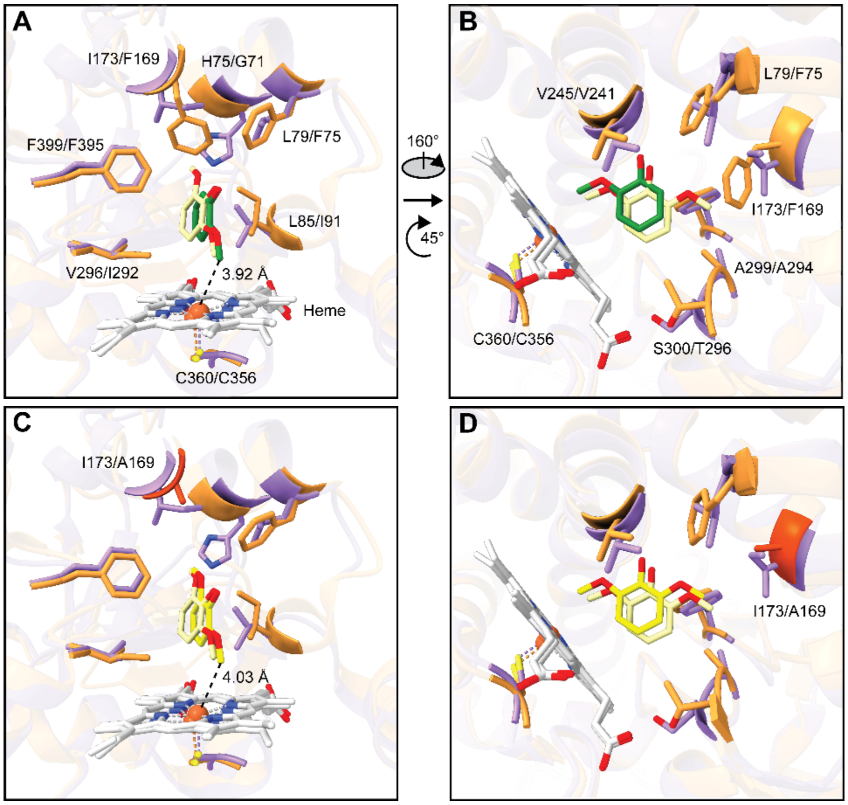


**Fig. S6.** Superposed structures of SyoA and GcoA; and SyoA and GcoA F169A highlighting differences in the active site residues. (A–B) Active site of SyoA bound to syringol (PDB: 8u19, purple) and GcoA bound to guaiacol (PDB: 5ncb, orange). Substrates are shown as stick diagrams with syringol in light yellow and guaiacol in green. (C–D) Active site of SyoA bound to syringol (PDB: 8u19, purple) and the GcoA mutant F169A bound to syringol (PDB:6qhh, orange). (C–D) Substrates shown as stick diagrams with syringol from SyoA in light yellow and syringol from GcoA F169A in dark yellow. F169A is highlighted in dark orange. The distance between the methoxy carbon and heme iron for the GcoA substrates is illustrated (black dashed line). The view of the active site in the right panel is made by rotating the top view 160° clockwise out of the plane and 45° clockwise in the plane.

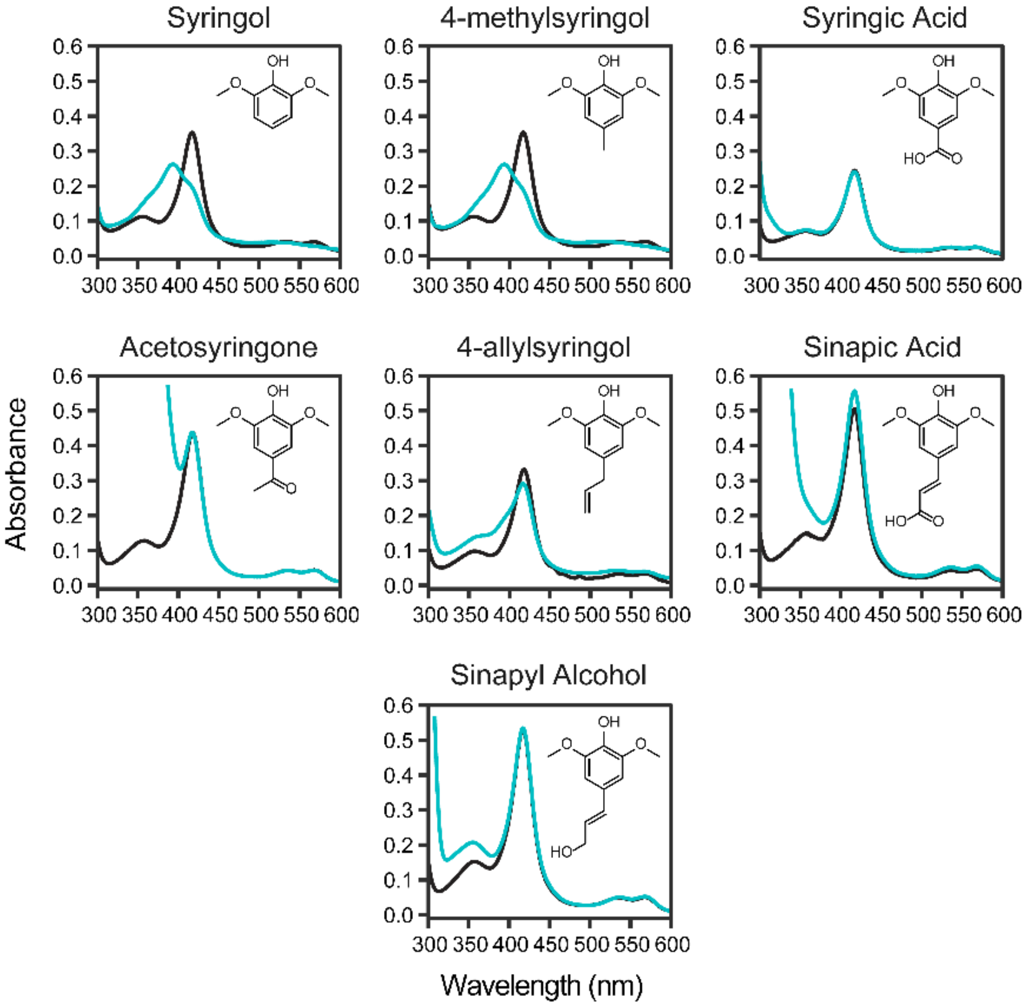


**Fig. S7.** Binding analysis of S-lignin monoaromatics to SyoA. *In vitro* spin-state shifts showing the Soret peak before (black) and after (blue) addition of an excess of each compound. A micromolar excess of each compound was titrated into a solution containing SyoA and the UV-Vis spectra monitored until the Soret band stopped shifting.

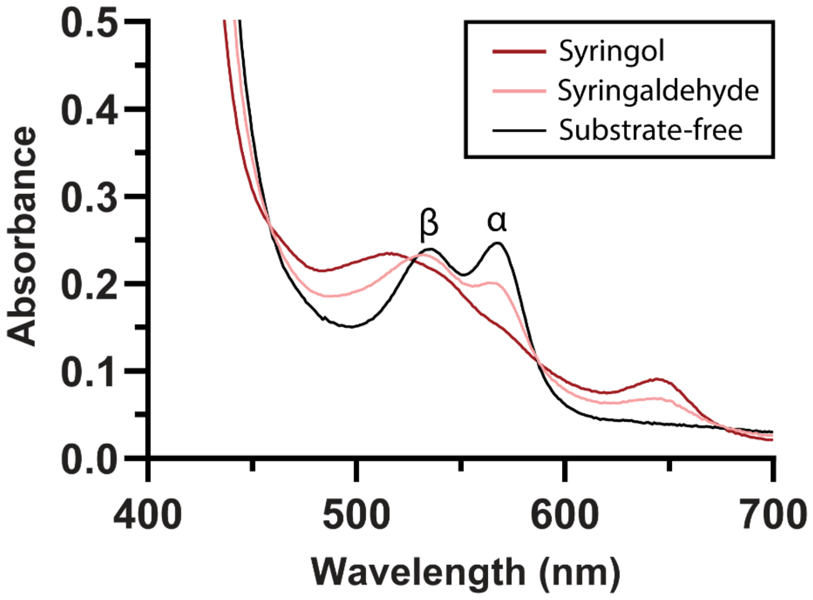


**Fig. S8.** Q-band analysis of syringaldehyde binding to SyoA using the α- and β-bands. In the absence of substrate, SyoA exhibits α- and β-bands at 568 and 535 nm, respectively. Addition of syringol (dark-red) results in merging of the bands at 515 nm and appearance of a charge transfer band at 640 nm. Addition of syringaldehyde (pink) results in a decrease in the α- and β-bands and appearance of a charge transfer band indicating binding of syringaldehyde.

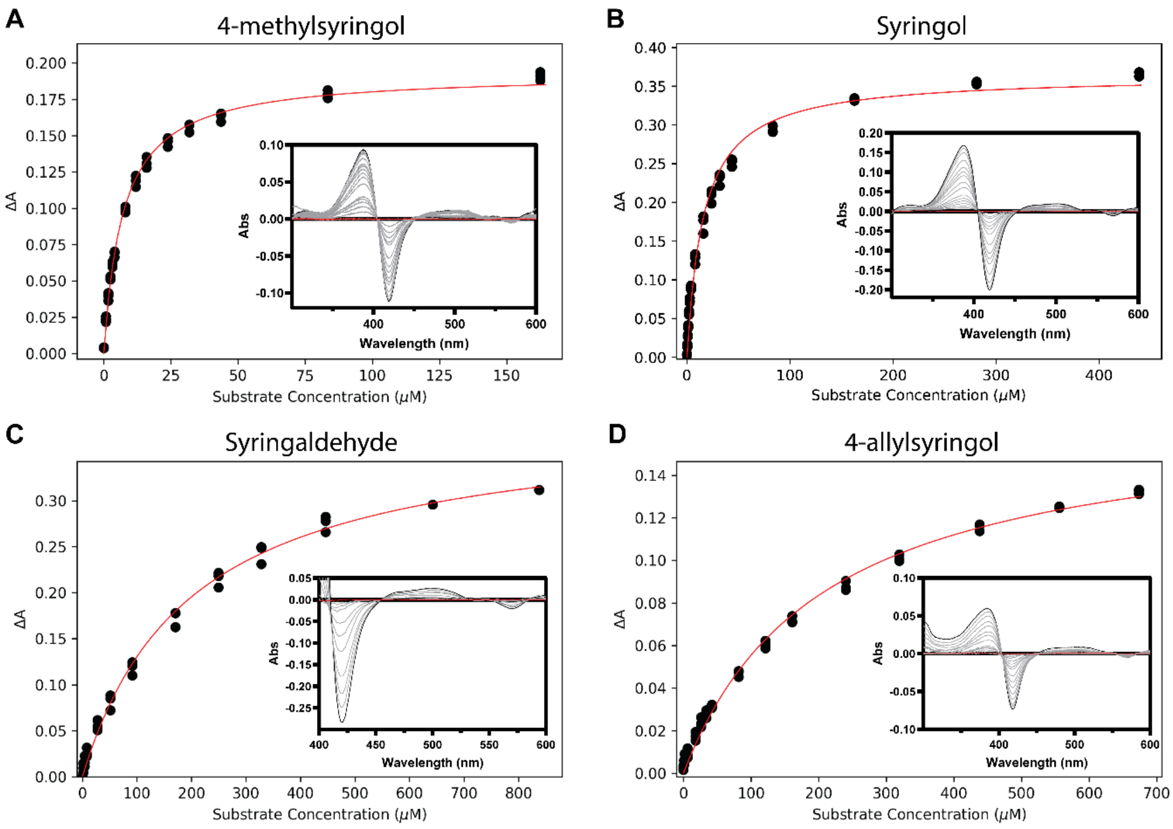


**Fig. S9.** Binding affinity curves of (A) 4-methylsyringol, (B) syringol, (C) syringaldehyde and (D) 4-allylsyringol to SyoA. The $K_D$ for 4-methylsyringol, syringol and 4-allylsyringol was determined by plotting the change in absorbance (ΔA) from the change in the peak (390 nm) to trough (419 nm). The $K_D$ for syringaldehyde was determined by plotting the change in absorbance (ΔA) at 419 nm due to strong absorbance from the substrate at 390 nm.

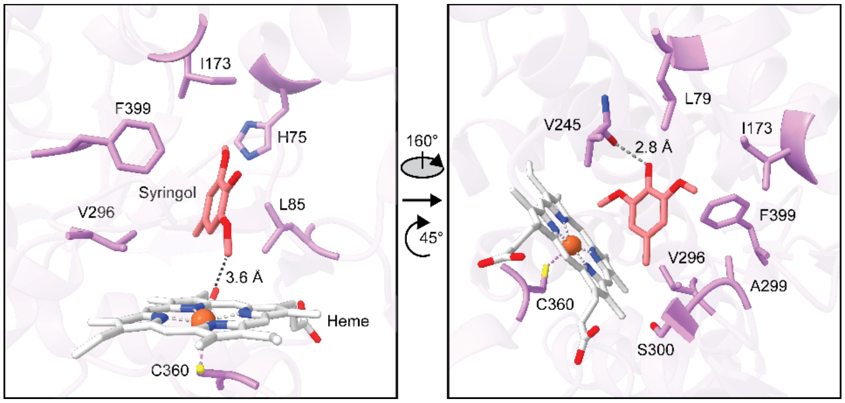


**Fig. S10.** Active site of SyoA bound to 4-methylsyringol (PDB: 8u1i) showing the key active site residues, highlighting the distance of the 2-methoxy group of 4-methylsyringol from the iron (left); and a side view highlighting the hydrogen bond distance between the 1-hydroxy group of syringol to the carbonyl of Val245 (right). The view of the active site from the side is made by rotating the top view 160° clockwise out of the plane and 45° clockwise in the plane.

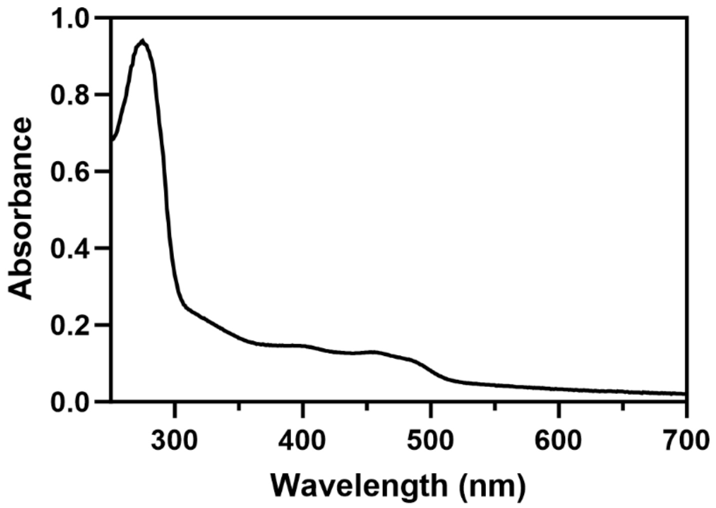


**Fig. S11.** UV-vis spectra of SyoB from *Amycolatopsis thermoflava* N1165. Spectra was measured in 50 mM Tris buffer pH 7.5. The gene encoding SyoA (Genbank: WP_037322545.1) is adjacent to its predicted redox partner SyoB (Genbank: WP_027930991). SyoB is a three-domain protein, with an N-terminal FMN-binding domain, followed by an NADH-binding domain and a 2Fe-2S domain. The UV-Vis spectrum of SyoB is typical for FMN and 2Fe-2S binding domains.

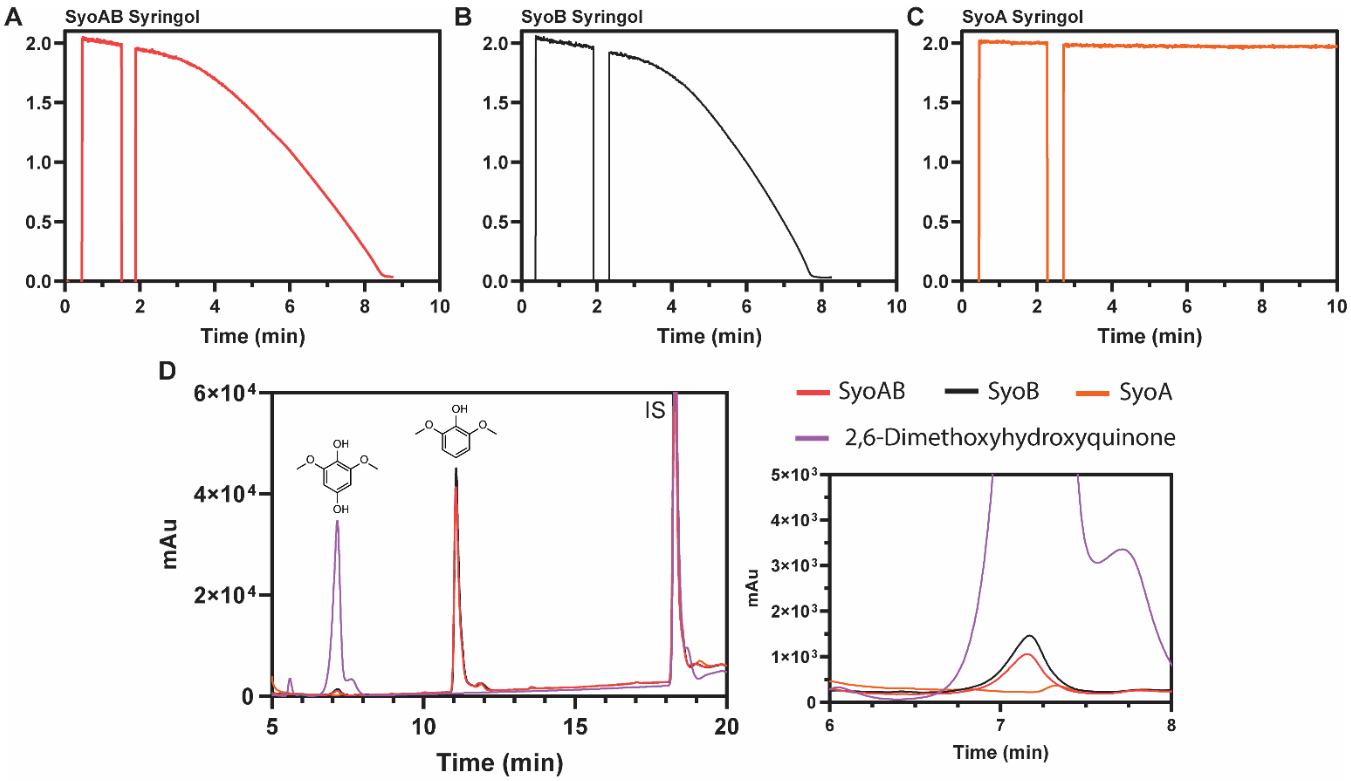


**Fig. S12.** *In vitro* NADH oxidation reactions of SyoAB with syringol. UV-Vis analysis of NADH consumption (340 nm) for reactions with syringol containing (A) SyoAB (B) SyoB and (C) SyoA. (D) HPLC chromatograms showing the formation of 2,6-dimethoxyhydroxyquinone in the presence of SyoB. Gradient: 20–95% MeCN in $H_2O$ with 0.1% v/v TFA. Detection wavelength: 254 nm. The absence of the expected product (3-methoxycatechol), formation of 2,6-dimethoxyhydroxyquinone and consumption of NADH in the negative control (SyoB) indicates uncoupling of electron transfer from the P450 activities. The observations above are consistent with syringol being reduced by SyoB in the presence of NADH and the organic product reacting with $O_2$ to generate superoxide and subsequently, hydrogen peroxide. The reactive superoxide could react with syringol to generate low levels of the para-quinone/para-hydroxyquinol (panels D and E). These species can be reduced more readily than syringol by SyoB, resulting in accelerating NADH consumption (panels A and B) and futile redox cycling between the quinone/hydroquinone forms, generating hydrogen peroxide; rather than channeling electrons to the P450 enzyme for substrate oxidation (6)

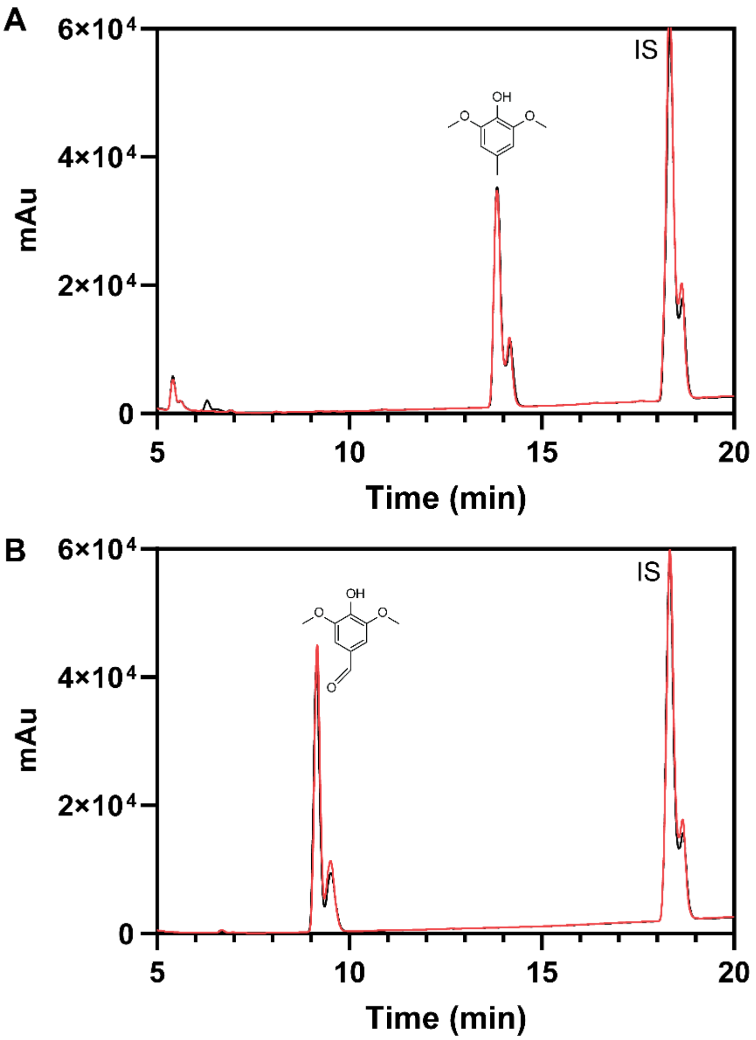


**Fig. S13.** HPLC chromatograms showing the *in vitro* NADH oxidation reactions of SyoAB with (A) 4-methylsyringol and (B) syringaldehyde. The control reactions without P450 are shown in black and the reaction with P450 in red. Gradient: 20–95% MeCN in $H_2O$ with 0.1% v/v TFA. Detection wavelength: 254 nm.

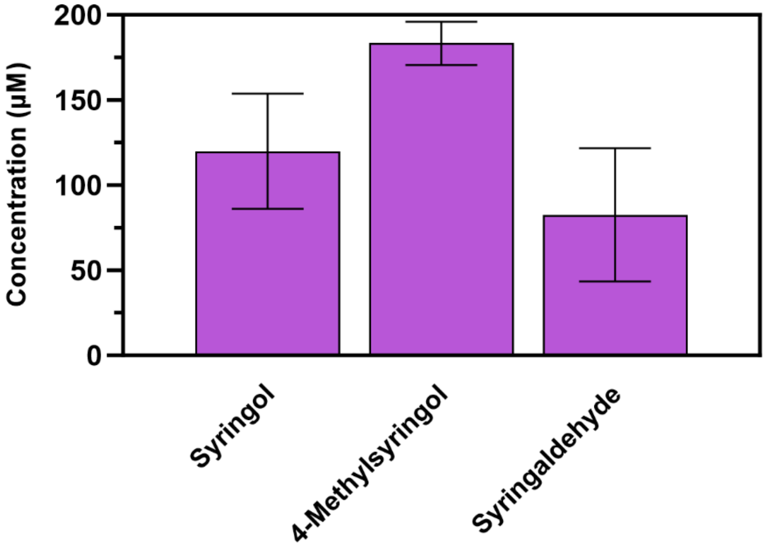


**Fig. S14.** Formaldehyde production during the $H_2O_2$-driven *O*-demethylation reactions with SyoA. Reactions containing 1 µM SyoA, 0.5 mM substrate and 10 mM $H_2O_2$ were quenched with catalase after 60 min and the concentration of formaldehyde estimated using the colorimetric Purpald assay. Formaldehyde is unstable in the presence of $H_2O_2$ resulting in an underestimate of the true concentration.

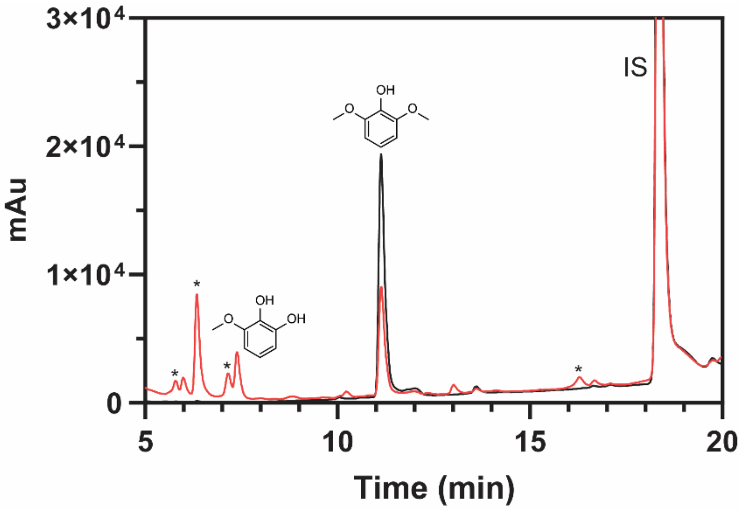


**Fig. S15**. HPLC chromatogram showing the $H_2O_2$-driven reaction of SyoA with syringol over 60 min. The control reaction without P450 is shown in black and the reaction with P450 in red. Syringol eluted at 11.2 min and 3-methoxycatechol 7.4 min. Gradient: 20–95% MeCN in $H_2O$ with 0.1% v/v TFA. Detection wavelength: 254 nm. *Denotes degradation products of 3-methoxycatechol by hydrogen peroxide.(2)

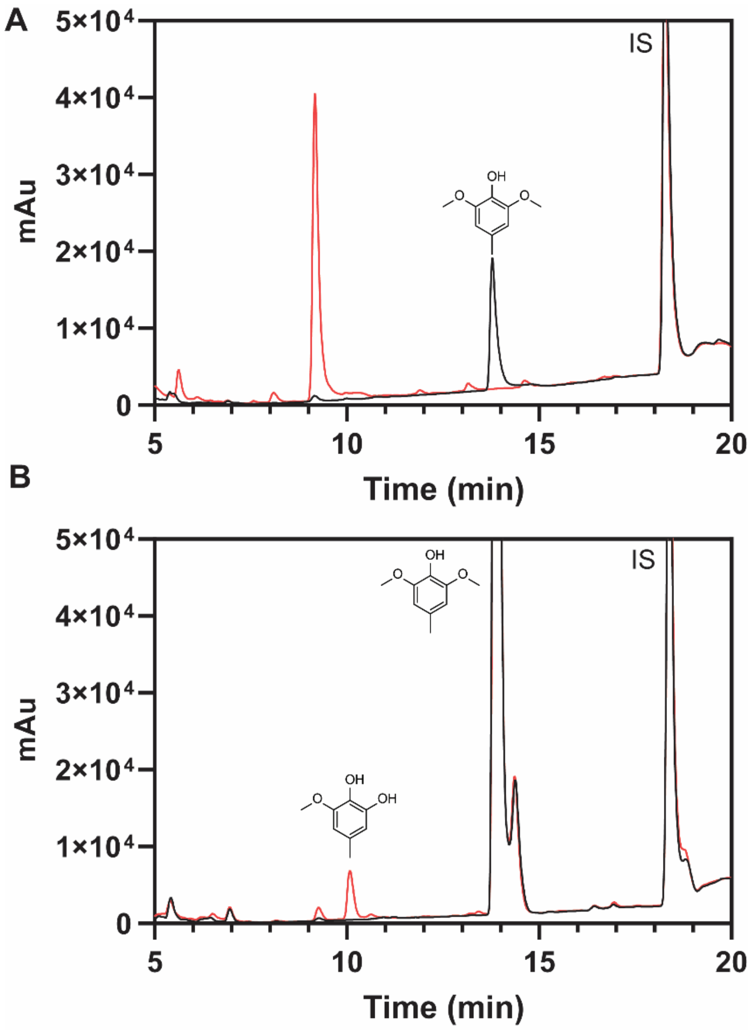


**Fig. S16.** HPLC chromatograms showing the $H_2O_2$-driven reaction of SyoA with 4-methylsyringol. Reactions were carried out using (A) 500 µM substrate and 10 mM $H_2O_2$ or (B) 5 mM substrate and 4 mM $H_2O_2$ over 60 mins. The control reactions without P450 are shown in black and the reaction with P450 in red. 4-methylsyringol eluted at 13.9 min, the unknown oxidation product in (A) at 9.1 min and the demethylated product (3-methoxy-5-methylbenzene-1,2-diol) in (B) at 10.1 min. The concentration of 3-methoxy-5-methylbenzene-1,2-diol was estimated using the calibration curve of 3-methoxycatechol due to both having a similar absorbance intensity at 254 nm (the wavelength used for HPLC analysis). Gradient: 20–95% MeCN in $H_2O$ with 0.1% v/v TFA. Detection wavelength: 254 nm.

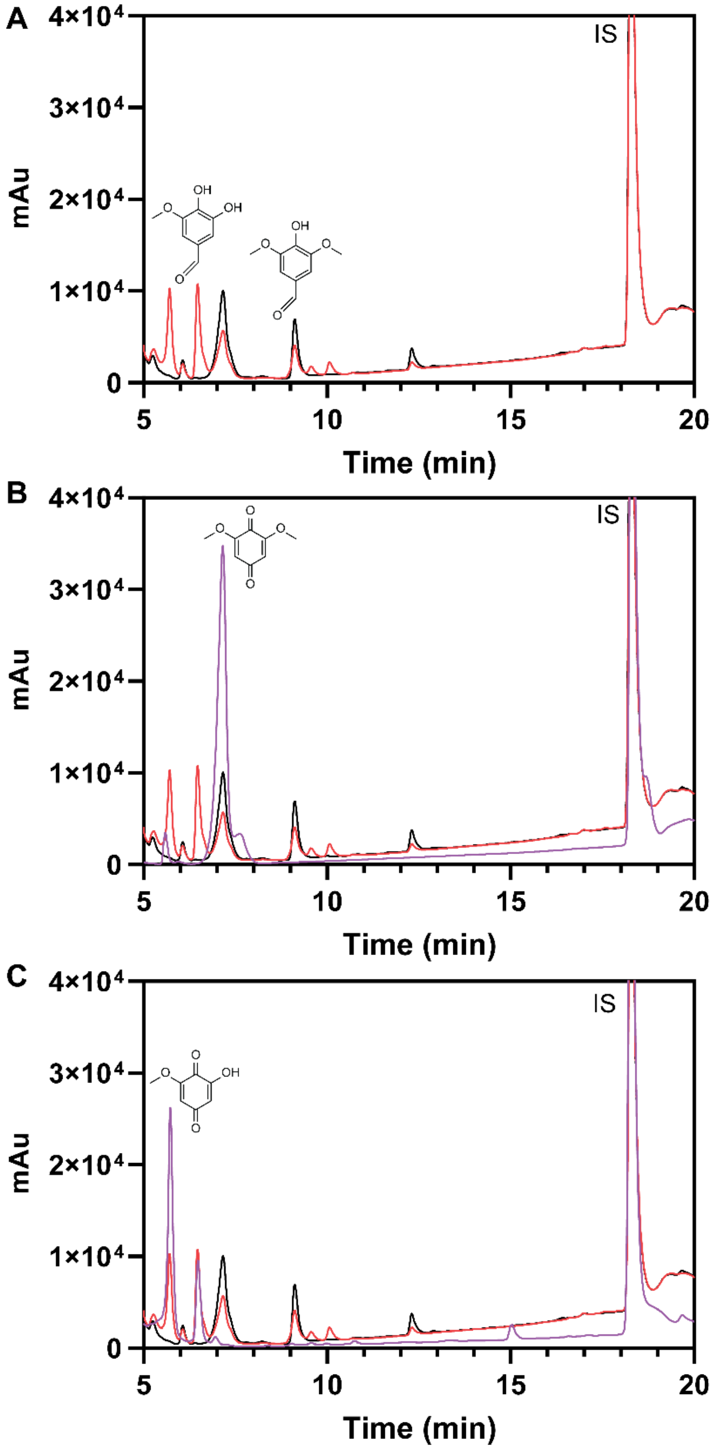


**Fig. S17.** HPLC chromatograms showing the (A) $H_2O_2$-driven reaction of SyoA with syringaldehyde. The control reactions without P450 are shown in black and the reaction with P450 in red. Syringaldehyde eluted at 9.1 min and the demethylated product (5-hydroxyvanillin) at 6.5 min. (B) Reaction compared to an authentic sample of 2,6-Dimethoxy-1,4-benzoquinone (purple) with elution time of 7.2 min. (C) Oxidation of 5-hydroxyvanillin in the presence of $H_2O_2$ (purple). The Dakin oxidation product of 5-hydroxyvanillin (2-hydroxy-6-methoxy-1,4-benzoquinone) has an elution time of 5.7 min. Gradient: 20–95% MeCN in $H_2O$ with 0.1% v/v TFA. Detection wavelength: 254 nm.

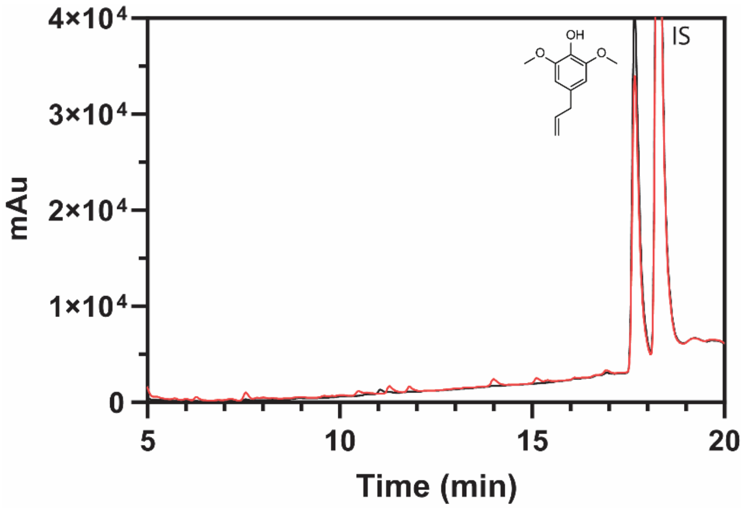


**Fig. S18.** HPLC chromatogram showing the $H_2O_2$-driven reaction of SyoA with 4-allylsyringol over 60 min. The control reaction without P450 is shown in black and the reaction with P450 in red. The demethylated products of syringol and 4-methylsyringol both had a retention time 3.7 min shorter than the respective substrate. A product with a retention time 3.7 min faster than 4-allylsyringol was also observed, which may indicate minor demethylation of 4-allylsyringol by SyoA. However, this product could not be confidently identified using a standard (not commercially available) or GC-MS. Gradient: 20–95% MeCN in $H_2O$ with 0.1% v/v TFA. Detection wavelength: 254 nm.

**A**

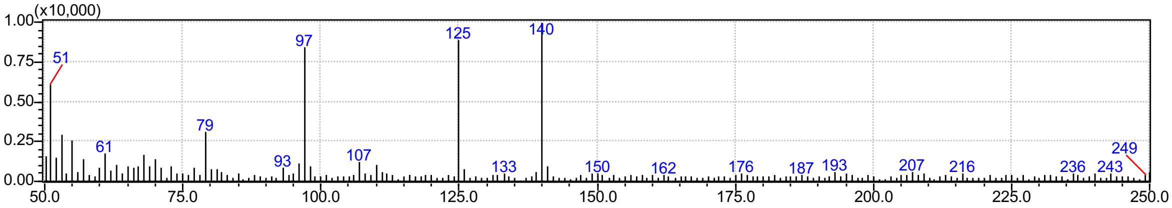


3-Methoxycatechol. RT 5.43 to 5.47 min. Observed m/z = 140. Expected m/z = 140.

**B**

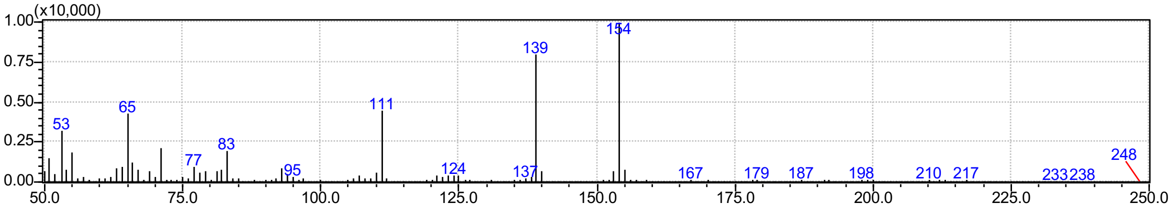


3-Methoxy-5-methylbenzene-1,2-diol. RT 6.75 to 6.80 min. Observed m/z = 154. Expected m/z = 154.

**C**

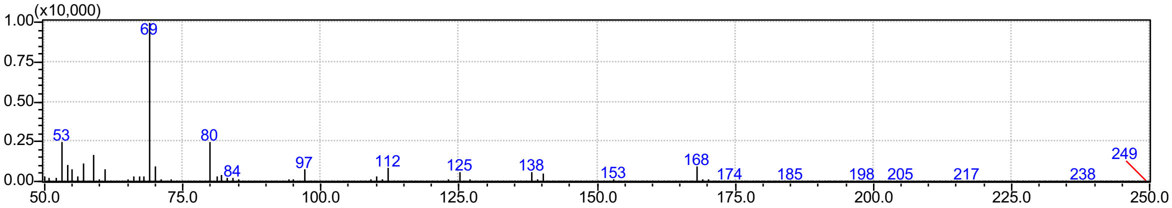


2,6-Dimethoxy-1,4-benzoquinone. RT 9.37 to 9.43 min. Observed m/z = 168. Expected m/z = 168.

**D**

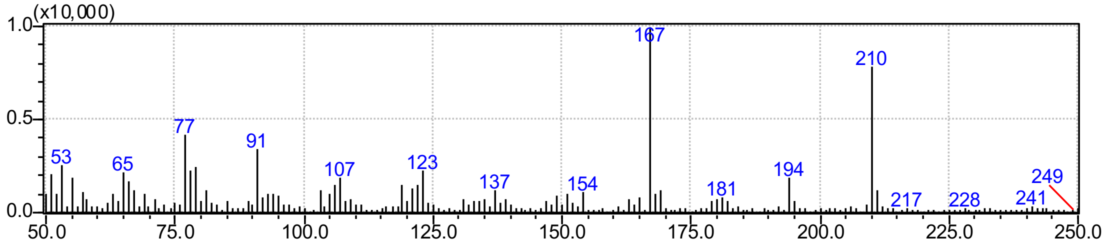


4-Allylsyringol +[O]. RT 13.72 to 13.82 min. Observed m/z = 210. Expected m/z = 210.

**Fig. S19.** Mass spectra of the major products identified from GC-MS analysis of the $H_2O_2$-driven reaction of SyoA with (A) Syringol (product: 3-methoxycatechol), (B) 4-methylsyringol (product: 3-methoxy-5-methylbenzene-1,2-diol), (C) syringaldehyde (product: 2,6-dimethoxy-1,4-benzoquinone) and (D) 4-allylsyringol (product: 4-Allylsyringol +[O]).

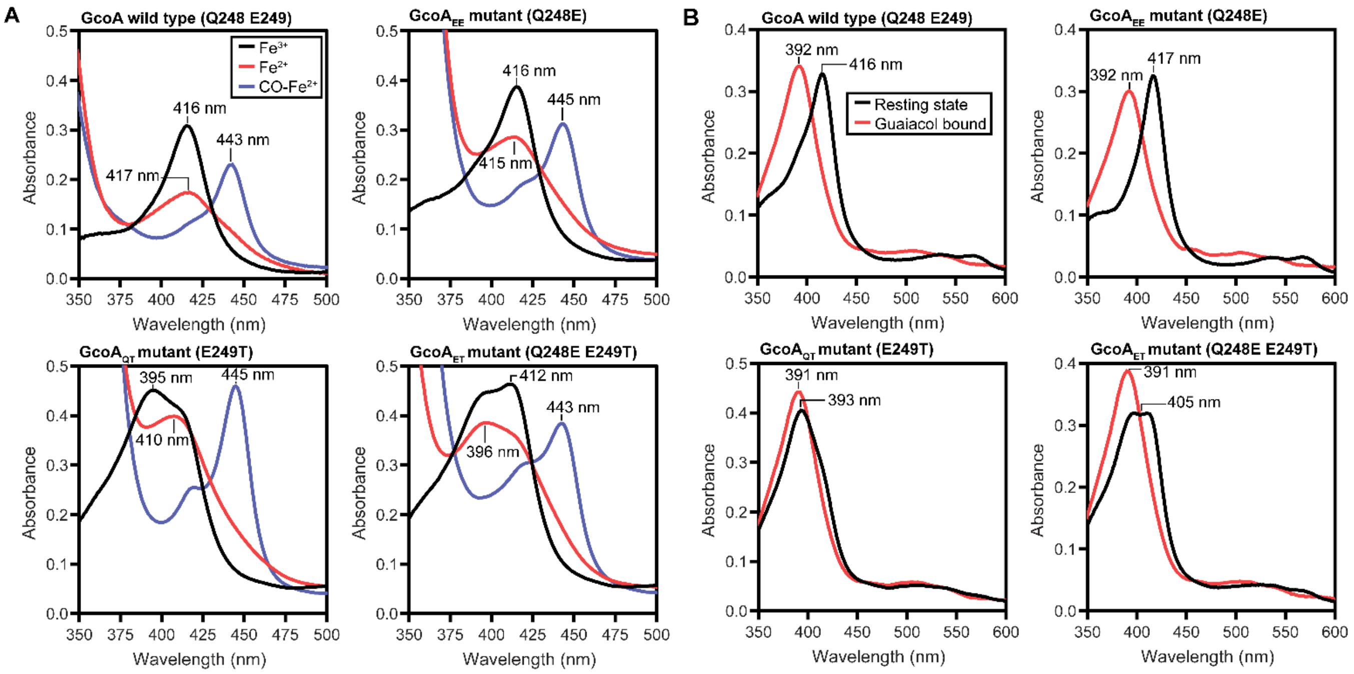


**Fig. S20.** Spectroscopic analysis showing the stability and substrate binding of the GcoA mutants. (A) UV-vis absorbance spectra of the ferric, ferrous and ferrous-CO bound state of GcoA mutants demonstrating proper folding and heme incorporation for all mutants (blue spectrum, Soret band at 443-445 nm). (B) UV-vis absorbance spectra demonstrating guaiacol binding to each mutant. GcoA mutants containing E249T exhibit blue-shifted Soret bands likely caused by interaction between threonine and the heme iron.

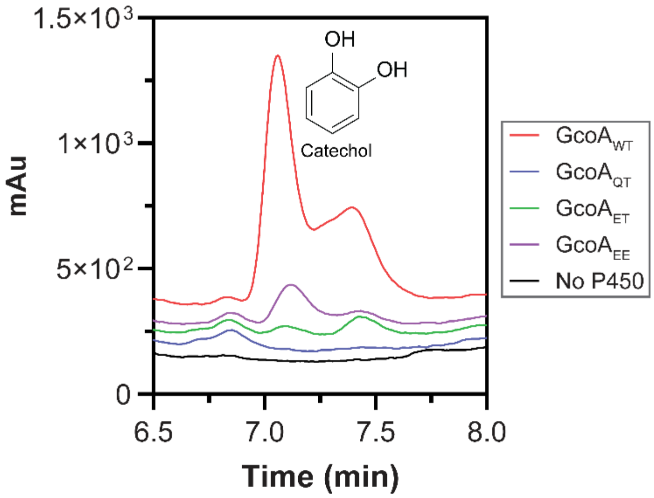


**Fig. S21.** Mutagenesis of the I-helix catalytic site residues Gln248 and Glu249 in GcoA cause a reduction in monooxygenase activity. HPLC chromatograms showing the amount of aromatic product (catechol) produced by each mutant compared to the wildtype. Reactions contained 0.5 μM GcoA, 1 μM GcoB, 320 μM NADH and 1 mM guaiacol. Gradient: 20–95% MeCN in $H_2O$ with 0.1% v/v TFA. Detection wavelength: 254 nm. Despite being folded, and competent for substrate binding, mutation of Gln248 and/or Glu249 resulted in a dramatic reduction in monooxygenase activity.

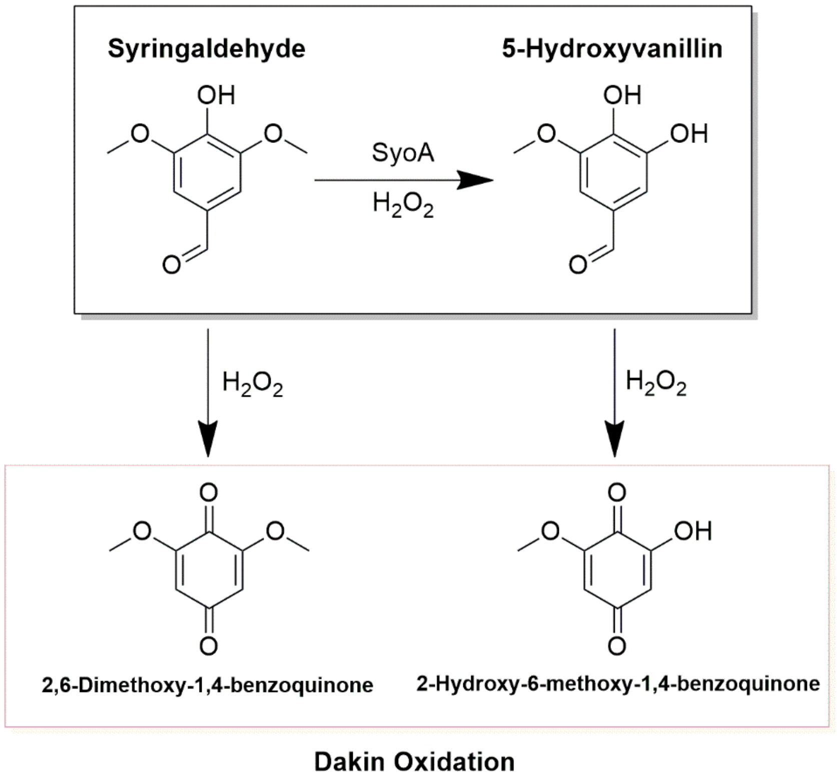


**Scheme S1.** Proposed Dakin oxidation products produced during the oxidation of syringaldehyde by SyoA.

**Table S1.** Crystallographic data statistics

| **Parameter** | **SyoA** | **SyoA with syringol** | **SyoA with 4-methylsyringol** |
|---|---|---|---|
| PDB code | 8u09 | 8u19 | 8u1i |
| Space group | $P\ 2_1\ 2_1\ 2$ | $P\ 2_1$ | $P\ 2_1$ |
| Wavelength (Å) | 0.9537 | 0.9537 | 0.9537 |
| Cell dimensions | | | |
| *a, b, c* (Å) | 54.86, 150.24, 60.64 | 57.85, 62.35, 58.71 | 57.41, 62.76, 58.08 |
| $\beta$ (°) | 90 | 103.06 | 104.58 |
| Resolution range (Å) | 47.19-1.98 (2.05-1.98) | 42.15-1.26 (1.31-1.26) | 41.60-1.12 (1.16-1.12) |
| $R_{merge}$ | 0.117 (0.631) | 0.050 (0.814) | 0.070 (1.213) |
| $R_{meas}$ | 0.122 (0.655) | 0.054 (0.882) | 0.075 (1.326) |
| $R_{pim}$ | 0.033 (0.175) | 0.021 (0.336) | 0.029 (0.527) |
| $CC_{1/2}$ | 0.999 (0.939) | 0.999 (0.818) | 0.999 (0.626) |
| Mean (i)/σ(i) | 19.84 (4.10) | 17.72 (1.90) | 11.41 (1.30) |
| Completeness (%) | 99.83 (98.75) | 98.70 (93.60) | 99.95 (99.93) |
| Multiplicity | 13.5 (13.7) | 6.6 (6.5) | 6.7 (6.1) |
| Structure refinement | | | |
| Resolution range (Å) | 47.19-1.98 | 42.15-1.26 | 41.60-1.12 |
| Unique reflections | 35629 | 108157 | 152917 |
| $R_{work}$ | 0.172 | 0.146 | 0.137 |
| $R_{free}$ | 0.205 | 0.182 | 0.153 |
| Total number of | | | |
| Non-hydrogen atoms | 3511 | 3916 | 3818 |
| Protein atoms | 3140 | 3330 | 3305 |
| Ligand atoms | 77 | 71 | 67 |
| Water molecules | 294 | 515 | 446 |
| RMSD | | | |
| Bond length (Å) | 0.005 | 0.006 | 0.008 |
| Bond angle (°) | 0.85 | 1.05 | 1.14 |

**Table S2.** Average RMSD of $C_\alpha$ atoms between the open and closed state of SyoA

| Helix/Loop | Average RMSD of $C_\alpha$ atoms (Å) |
|---|---|
| B'-Helix | 3.0 |
| F-Helix | 4.0 |
| F/G-loop | 8.3 |
| G-Helix | 6.2 |
| H-Helix | 3.3 |
| H/I-loop | 4.2 |
| I-Helix | 1.2 |

**Table S3.** Relative disposition of the $C_α$ atoms for the active site residues between the open and closed state of SyoA

| Residue | RMSD (Å) |
|---|---|
| H75 | 2.5 |
| L79 | 2.6 |
| L85 | 0.9 |
| I173 | 4.4 |
| V245 | 1.5 |
| L248 | 1.6 |
| E253 | 1.2 |
| V296 | 0.6 |
| A299 | 0.6 |
| S300 | 0.7 |
| F399 | 0.8 |

**Table S4.** Heme geometry. No obvious heme ruffling or doming was observed in the substrate-bound structure, with the heme appearing to remain planar in both open and closed states. Minor changes were observed in the position of the heme iron. The iron moved ~0.2 Å out of the heme plane in a concerted motion with the axial cysteine (Cys360). This movement resulted in an increase in the average ∠N−Fe−S angle by ~5.0°.

| **Distance (Å)** | **Structure** | | |
|---|---|---|---|
| | **Substrate-free** | **Syringol-bound** | **4-methylsyringol-bound** |
| **Fe−OH₂** | 2.48 | NA | NA |
| **Fe−S** | 2.24 | 2.30 | 2.31 |
| **Fe−NA** | 2.07 | 2.00 | 2.04 |
| **Fe−NB** | 2.04 | 2.05 | 2.03 |
| **Fe−NC** | 2.05 | 2.03 | 2.03 |
| **Fe−ND** | 2.03 | 2.05 | 2.05 |
| **Angle (°)** | | | |
| **∠NA−Fe−S** | 97.8 | 102.0 | 101.2 |
| **∠NB−Fe−S** | 89.0 | 93.3 | 93.6 |
| **∠NC−Fe−S** | 86.8 | 92.9 | 94.0 |
| **∠ND−Fe−S** | 97.1 | 103.9 | 103.6 |

**Table S5.** DNA and amino acid sequences for the proteins used in this study. Restriction sites (underlined), stop codon (italics) and affinity tags (bold).

| | | |
|---|---|---|
| **SyoA pET3a** | **DNA Sequence** | **ATGGGTGGCTCCCATCACCATCACCATCACGACTACGATATCCCGACCACCGAAAACCTGTACTTCCAGGGC**<br>**GCC**ATGACAACAAAGCATACTACAGCCGGCGATACCCAAGAATGGTTAGCGACTGTAACCGTAGAGCAGTTA<br>GAGAATGACCCATACCCTATCTTTGAGCGTTTACGCCGCGAGGCCCCAGTCGCCTGGATCCCGGCCGCCCAT<br>GCCTGGGTAGCATCTACGTGGGAAGCCTGTCGTACGATTGCCGACGATGCAACTAACTTTCGCGGTGGAACT<br>TCACCTATGCACGAGCGCGTACTTGGCACGGATCACATCCTGGGAGCCGAGGGTGAGACGCACCAAGACCTT<br>CGCGCAGCCGTAGACCCGCCTCTTAAACCCCGTGCGTTCCGCCCCTTGCTTGAGGAGCAGGTACGTCCAACG<br>GTCCGTCGTTACCTTGAGGCAATTCGCGGACAGGGCAAGGCAGAACTGATGGCTGACTACTTTGAGCCGATT<br>TCTGTGCGTTGCGTGGGTGACGTCATTGGGCTTACGGACGTGGACAGTGACACGTTGCGCCGCTGGTTTCAT<br>GCATTAGCGCGTGGAATCGCAAACACGGCAATGGACGCGGAAGGTCGTTTCACTAATCCGGGCGGCTTTGCC<br>CCTGCGGATGAGGCAGGTGCCGAAATTCGCGAAGTATTAGAGCCGCTCGTAGCCAAGTTATCTGCTGAGCCG<br>GACGGCAGTGCGCTTAGCCACTACCTTCATCATGGACGCCCTCATGGCGATCCTCGTACACTTGAGCAACTG<br>TTACCGTCCCTCAAGGTGATCATTCTCGGCGGGCTGCAGGAACCGGGTCACCAATGTGCAGCCACCTTCCTG<br>GGTCTCACAACTCGTCCTGAACAGTTAAAGCGCGTGACCGAAGACGCCACGCTCCTGCCACGCGCGCTCACT<br>GAAGGACTCCGCTGGATGAGCCCCGTATTCTCCGCGAGCAGTCGTTTGCCTTTACGCGAGATTACGATGGGA<br>GAAGCGACGATGCGTCCGGGCCAGACGGTTTGGTTGAGTTATGGGAGTGCAAACCGCGACGAGGCCGTGTTT<br>GACCGTCCGGATGTATTCGATCTTGATCGTGCCACTCATCCACATCTGGCGTTCGGCACCGGTCGCCACCTT<br>TGTAGTGGTTCAGCCTATGCCCCACAGGTGGCCCGTATTGCGCTCGAAGAGCTGTTTACAGCCTTTCCCAGC<br>ATTCGTCTTGATCCAGCGCATGAAGTACCTGTGTGGGGCTGGCTGTTCCGCGGGCCTCAACGCCTGGATGTG<br>CTGTGGGAC*TGA* |
| | **Amino Acid Sequence** | **MGGSHHHHHHDYDIPTTENLYFQGA**MTTKHTTAGDTQEWLATVTVEQLENDPYPIFERLRREAPVAWIPAAH<br>AWVASTWEACRTIADDATNFRGGTSPMHERVLGTDHILGAEGETHQDLRAAVDPPLKPRAFRPLLEEQVRPT<br>VRRYLEAIRGQGKAELMADYFEPISVRCVGDVIGLTDVDSDTLRRWFHALARGIANTAMDAEGRFTNPGGFA<br>PADEAGAEIREVLEPLVAKLSAEPDGSALSHYLHHGRPHGDPRTLEQLLPSLKVIILGGLQEPGHQCAATFL<br>GLTTRPEQLKRVTEDATLLPRALTEGLRWMSPVFSASSRLPLREITMGEATMRPGQTVWLSYGSANRDEAVF<br>DRPDVFDLDRATHPHLAFGTGRHLCSGSAYAPQVARIALEELFTAFPSIRLDPAHEVPVWGWLFRGPQRLDV<br>LWD |
| **SyoB gBlock** | **DNA Sequence** | ATTTTACATATGACGCGTGAGCTGGTGGTCACATCTAAGGAGGAAGCAGCTGAGGATGTTATGGTTATCCAC<br>CTGACGGACCCCGGAGGCGATCCACTGCCCGAATGGACCCCCGGAGCGCACGTAGGTGTCGATGTGGACGGG<br>GATGTACGCCAGTACTCACTTTGCGGACCCACTAGCGAACGTTTCACCTGGCGCATCGGCGTCCGTCGTGAA<br>TCCAACGGAGTCGTATCGAACAAATTGCACGACAGTGTGTCGGTAGGGCGTCGTCTGCGCGTTAGTGACCCG<br>GAAAACTTATTCCCATTGGTTCCCGCCGAACGCTATGTATTTATTGCCGGAGGTATCGGGATTACCCCGCTT<br>TTGCCCATGCTTGAAGAAGTCAAACGTGCCGGTCGCGAGTGGAGTCTTTATTACGGCGGGCGCAGTCGCCGT<br>CACATGGCTTTCTGCTCCGAGGTTGATGGTCCTGGAGTTACTTTATGGCCTGAGGACGAACACGGCTTACTG<br>CCCGTAAATTCACTTCTGGGAGAGCCACGTCCTGGTACTGCTGTGTACTGCTGTGGGCCAGCTCCCTTAATC<br>GATGCGGTCACGGAGGCTTGTGCCGCATGGCCCGCGGGTACTTTGCATGTCGAGCGCTTCACCCCCGTCCGC<br>CCAGGGGACAGCGCGCGTCCTTTTGTTGCTGAACTTCGCCGCTCCAATCGCACCATCGAAGTGCCCGCCGAC<br>AAAAGTCTTTTAGAGGCAGTCGAAGCAGCCGGAATTCCCGTCCTGTCCAGCTGTCGTACGGGTACATGTGGC<br>ACTTGCGAGGCGACCGTCCTGGACGGTGAGCCCGAGCACCATGATGAGGTTCTTACGGATGAGGAGCGTCAG<br>GACGGGAAGTTAATTATGTTATGTGTCTCCCGTAGTCGCTCAGCGGTGCTGGGGTTGGATCTT*TAATAG*GGT<br>ACCAAGCTTTTAATT |
| | **Amino Acid Sequence** | MTRELVVTSKEEAAEDVMVIHLTDPGGDPLPEWTPGAHVGVDVDGDVRQYSLCGPTSERFTWRIGVRRESNG<br>VVSNKLHDSVSVGRRLRVSDPENLFPLVPAERYVFIAGGIGITPLLPMLEEVKRAGREWSLYYGGRSRRHMA<br>FCSEVDGPGVTLWPEDEHGLLPVNSLLGEPRPGTAVYCCGPAPLIDAVTEACAAWPAGTLHVERFTPVRPGD<br>SARPFVAELRRSNRTIEVPADKSLLEAVEAAGIPVLSSCRTGTCGTCEATVLDGEPEHHDEVLTDEERQDGK<br>LIMLCVSRSRSAVLGLDL |
| **SyoB pET29b** | **DNA Sequence** | CATATGACGCGTGAGCTGGTGGTCACATCTAAGGAGGAAGCAGCTGAGGATGTTATGGTTATCCACCTGACG<br>GACCCCGGAGGCGATCCACTGCCCGAATGGACCCCCGGAGCGCACGTAGGTGTCGATGTGGACGGGGATGTA<br>CGCCAGTACTCACTTTGCGGACCCACTAGCGAACGTTTCACCTGGCGCATCGGCGTCCGTCGTGAATCCAAC |

| | | |
|---|---|---|
| | | GGAGTCGTATCGAACAAATTGCACGACAGTGTGTCGGTAGGGCGTCGTCTGCGCGTTAGTGACCCGGAAAAC<br>TTATTCCCATTGGTTCCCGCCGAACGCTATGTATTTATTGCCGGAGGTATCGGGATTACCCCGCTTTTGCCC<br>ATGCTTGAAGAAGTCAAACGTGCCGGTCGCGAGTGGAGTCTTTATTACGGCGGGCGCAGTCGCCGTCACATG<br>GCTTTCTGCTCCGAGGTTGATGGTCCTGGAGTTACTTTATGGCCTGAGGACGAACACGGCTTACTGCCCGTA<br>AATTCACTTCTGGGAGAGCCACGTCCTGGTACTGCTGTGTACTGCTGTGGGCCAGCTCCCTTAATCGATGCG<br>GTCACGGAGGCTTGTGCCGCATGGCCCGCGGGTACTTTGCATGTCGAGCGCTTCACCCCCGTCCGCCCAGGG<br>GACAGCGCGCGTCCTTTTGTTGCTGAACTTCGCCGCTCCAATCGCACCATCGAAGTGCCCGCCGACAAAAGT<br>CTTTTAGAGGCAGTCGAAGCAGCCGGAATTCCCGTCCTGTCCAGCTGTCGTACGGGTACATGTGGCACTTGC<br>GAGGCGACCGTCCTGGACGGTGAGCCCGAGCACCATGATGAGGTTCTTACGGATGAGGAGCGTCAGGACGGG<br>AAGTTAATTATGTTATGTGTCTCCCGTAGTCGCTCAGCGGTGCTGGGGTTGGATCTT**CACCACCACCACCAC**<br>**CAC***TAATAG*AAGCTT |
| | **Amino Acid Sequence** | MTRELVVTSKEEAAEDVMVIHLTDPGGDPLPEWTPGAHVGVDVDGDVRQYSLCGPTSERFTWRIGVRRESNG<br>VVSNKLHDSVSVGRRLRVSDPENLFPLVPAERYVFIAGGIGITPLLPMLEEVKRAGREWSLYYGGRSRRHMA<br>FCSEVDGPGVTLWPEDEHGLLPVNSLLGEPRPGTAVYCCGPAPLIDAVTEACAAWPAGTLHVERFTPVRPGD<br>SARPFVAELRRSNRTIEVPADKSLLEAVEAAGIPVLSSCRTGTCGTCEATVLDGEPEHHDEVLTDEERQDGK<br>LIMLCVSRSRSAVLGLDL**HHHHHH** |
| **GcoA$_{EE}$ Mutant pET28a** | **DNA Sequence** | CC**ATGGGTGGCTCCAGCCATCACCATCACCATCACAGCAGCGGCGAAAACCTGTACTTCCAGGGCCAT**ATGA<br>CCACAACAGAAAGACCGGACCTTGCATGGCTCGACGAGGTAACAATGACCCAGTTAGAAAGAAACCCTTATG<br>AGGTATATGAGAGACTGCGCGCTGAGGCGCCGCTGGCTTTTGTCCCAGTTCTGGGCTCCTATGTTGCAAGTA<br>CTGCGGAAGTCTGCCGAGAAGTAGCCACAAGCCCAGATTTCGAAGCAGTGATTACTCCGGCCGGCGGTCGTA<br>CCTTTGGACACCCGGCAATCATTGGTGTGAATGGGGATATCCATGCGGATTTGCGTTCAATGGTTGAACCTG<br>CCTTACAGCCGGCAGAGGTGGACCGATGGATAGATGACCTGGTGCGGCCGATTGCACGTCGCTACCTTGAAA<br>GATTTGAAAATGATGGTCATGCAGAGCTTGTGGCACAGTACTGTGAACCTGTGAGTGTCCGTTCGTTGGGTG<br>ACCTCTTAGGCCTGCAAGAAGTGGATAGTGATAAACTGCGTGAATGGTTTGCTAAACTGAATCGGTCGTTCA<br>CGAATGCGGCCGTGGACGAAAATGGAGAATTTGCCAATCCCGAAGGTTTCGCTGAGGGCGACCAAGCAAAAG<br>CAGAGATCCGAGCAGTGGTGGATCCGCTGATAGATCGCTGGATAGAACATCCAGACGACAGCGCTATCTCCC<br>ATTGGTTACATGACGGAATGCCCCCTGGTCAGACACGCGATCGTGAGTACATATACCCGACCATTTACGTTT<br>ACTTGCTGGGTGCAATGGAAGAGCCTGGTCACGGTATGGCGTCGACCCTCGTAGGTCTGTTTTCAAGACCGG<br>AACAGTTAGAGGAAGTTGTTGACGACCCGACCTTGATACCTAGAGCCATTGCGGAAGGCCTTCGCTGGACCT<br>CGCCGATATGGTCCGCTACTGCGAGAATTAGCACCAAACCGGTAACTATAGCAGGGGTGGATCTTCCGGCGG<br>GCACTCCGGTCATGTTGTCTTACGGTTCGGCAAATCACGACACAGGCAAATACGAGGCACCTTCACAATACG<br>ACCTGCACAGACCGCCGCTGCCACATCTGGCCTTTGGTGCCGGTAACCACGCCTGCGCGGGGATATATTTCG<br>CTAACCATGTTATGCGGATCGCACTGGAAGAACTGTTTGAGGCAATCCCTAATCTCGAAAGAGATACGCGCG<br>AGGGTGTCGAATTCTGGGGTTGGGGTTTCCGTGGTCCCACAAGCCTGCACGTAACTTGGGAAGTT*TAATAG*G<br>GTACCAAGCTTCTCGAG |
| | **Amino Acid Sequence** | **MGGSSHHHHHHSSGENLYFQGH**MTTTERPDLAWLDEVTMTQLERNPYEVYERLRAEAPLAFVPVLGSYVAST<br>AEVCREVATSPDFEAVITPAGGRTFGHPAIIGVNGDIHADLRSMVEPALQPAEVDRWIDDLVRPIARRYLER<br>FENDGHAELVAQYCEPVSVRSLGDLLGLQEVDSDKLREWFAKLNRSFTNAAVDENGEFANPEGFAEGDQAKA<br>EIRAVVDPLIDRWIEHPDDSAISHWLHDGMPPGQTRDREYIYPTIYVYLLGAMEEPGHGMASTLVGLFSRPE<br>QLEEVVDDPTLIPRAIAEGLRWTSPIWSATARISTKPVTIAGVDLPAGTPVMLSYGSANHDTGKYEAPSQYD<br>LHRPPLPHLAFGAGNHACAGIYFANHVMRIALEELFEAIPNLERDTREGVEFWGWGFRGPTSLHVTWEV |
| **GcoA$_{QT}$ Mutant pET28a** | **DNA Sequence** | CC**ATGGGTGGCTCCAGCCATCACCATCACCATCACAGCAGCGGCGAAAACCTGTACTTCCAGGGCCAT**ATGA<br>CAACAACCGAACGTCCGGACCTTGCATGGCTGGACGAGGTGACCATGACCCAGTTAGAACGGAACCCGTATG<br>AGGTTTATGAGCGTCTGCGCGCCGAGGCGCCATTAGCTTTTGTTCCGGTGCTGGGCTCGTATGTTGCAAGCA<br>CAGCGGAAGTCTGCAGAGAAGTGGCTACCTCTCCAGATTTCGAAGCGGTCATTACTCCAGCAGGTGGTCGTA<br>CGTTTGGGCACCCTGCAATAATAGGAGTCAATGGAGATATACATGCCGATTTGCGGTCAATGGTCGAACCCG<br>CGCTGCAGCCAGCAGAGGTGGACCGATGGATAGATGACCTGGTTCGTCCGATAGCCCGGCGCTACTTGGAAC<br>GATTTGAAAATGATGGCCATGCAGAGCTGGTCGCACAGTACTGTGAACCTGTCTCAGTCCGTTCGCTGGGTG |

| | | |
|---|---|---|
| | | ACTTACTGGGTTTACAAGAAGTGGATAGTGATAAATTGAGAGAATGGTTTGCGAAACTTAATCGGAGCTTCA<br>CAAATGCGGCCGTGGACGAAAATGGCGAATTTGCTAATCCAGAAGGTTTCGCCGAGGGAGACCAAGCCAAAG<br>CGGAGATCCGGGCGGTAGTTGATCCGCTTATAGATCGGTGGATAGAACATCCGGACGACTCCGCTATCTCAC<br>ATTGGCTCCATGACGGCATGCCCCCTGGACAGACCCGCGATCGTGAGTACATCTACCCTACTATATACGTTT<br>ACTTACTGGGTGCCATGCAAACCCCTGGACACGGTATGGCCAGCACACTTGTTGGTCTGTTTTCACGCCCGG<br>AACAGTTAGAGGAAGTAGTGGACGACCCGACGCTCATTCCCCGGGCCATTGCCGAAGGCTTACGCTGGACTA<br>GTCCAATTTGGTCCGCGACGGCTCGAATTTCTACAAAACCCGTCACCATAGCCGGCGTGGATTTGCCAGCGG<br>GGACTCCTGTTATGCTTTCTTACGGGAGTGCGAATCACGACACAGGTAAATACGAGGCGCCGTCGCAATACG<br>ATCTCCACCGACCGCCCCTGCCTCATCTGGCATTTGGCGCAGGAAACCACGCTTGCGCCGGGATCTATTTCG<br>CGAACCATGTGATGCGGATCGCTCTTGAAGAACTGTTTGAGGCAATCCCAAATCTGGAGCGTGATACACGCG<br>AGGGCGTGGAATTCTGGGGATGGGGATTCCGAGGCCCGACATCCCTGCACGTAACCTGGGAAGTT*TAATAG*G<br>GTACCAAGCTTCTCGAG |
| | **Amino Acid Sequence** | **MGGSSHHHHHHSSGENLYFQGH**MTTTERPDLAWLDEVTMTQLERNPYEVYERLRAEAPLAFVPVLGSYVAST<br>AEVCREVATSPDFEAVITPAGGRTFGHPAIIGVNGDIHADLRSMVEPALQPAEVDRWIDDLVRPIARRYLER<br>FENDGHAELVAQYCEPVSVRSLGDLLGLQEVDSDKLREWFAKLNRSFTNAAVDENGEFANPEGFAEGDQAKA<br>EIRAVVDPLIDRWIEHPDDSAISHWLHDGMPPGQTRDREYIYPTIYVYLLGAMQTPGHGMASTLVGLFSRPE<br>QLEEVVDDPTLIPRAIAEGLRWTSPIWSATARISTKPVTIAGVDLPAGTPVMLSYGSANHDTGKYEAPSQYD<br>LHRPPLPHLAFGAGNHACAGIYFANHVMRIALEELFEAIPNLERDTREGVEFWGWGFRGPTSLHVTWEV |
| **GcoA$_{ET}$ Mutant pET28a** | **DNA Sequence** | CC**ATGGGTGGCTCCAGCCATCACCATCACCATCACAGCAGCGGCGAAAACCTGTACTTCCAGGGCCAT**ATGA<br>CAACAACTGAACGTCCGGACCTCGCATGGCTGGACGAGGTGACCATGACGCAGTTAGAACGGAACCCGTATG<br>AGGTCTATGAGAGATTACGGGCGGAAGCCCCGCTGGCGTTTGTGCCGGTTCTTGGCTCCTATGTTGCCAGCA<br>CGGCCGAAGTGTGCCGGGAAGTAGCCACATCACCTGATTTCGAAGCCGTGATTACTCCAGCAGGCGGTCGGA<br>CGTTTGGTCACCCGGCAATCATTGGGGTCAATGGGGATATCCATGCCGATCTGCGTTCAATGGTAGAACCGG<br>CACTGCAGCCGGCAGAGGTCGACCGGTGGATCGATGACTTAGTACGGCCGATTGCGCGGCGCTACCTCGAAC<br>GGTTTGAAAATGATGGCCATGCCGAGCTTGTCGCTCAGTACTGTGAACCTGTCAGCGTACGGTCACTGGGTG<br>ACTTGTTAGGTCTCCAAGAAGTAGATTCAGATAAACTGAGAGAATGGTTTGCTAAATTGAATCGGTCTTTCA<br>CGAATGCCGCAGTAGACGAAAATGGTGAATTTGCTAATCCTGAAGGTTTCGCCGAGGGCGACCAAGCTAAAG<br>CGGAGATCCGGGCAGTCGTTGATCCGCTTATCGATCGGTGGATAGAACATCCAGACGACTCTGCCATCTCCC<br>ATTGGTTGCATGACGGCATGCCGCCTGGGCAGACTCGCGATCGGGAGTACATCTACCCTACCATTTACGTTT<br>ACTTACTGGGTGCTATGGAGACACCCGGTCACGGGATGGCTTCTACACTTGTTGGTTTGTTTTCACGCCCAG<br>AACAGCTGGAAGAAGTTGTAGACGACCCCACATTGATTCCACGTGCTATAGCTGAAGGATTGCGCTGGACTT<br>CACCGATTTGGTCAGCTACGGCACGAATTAGTACTAAACCCGTCACGATCGCCGGCGTGGATCTTCCGGCAG<br>GAACCCCGGTAATGTTGTCCTACGGCTCAGCAAATCACGACACTGGCAAATACGAGGCACCGTCTCAATACG<br>ACTTACACCGACCGCCGTTACCTCATCTGGCCTTTGGTGCAGGTAACCACGCATGCGCCGGAATATATTTCG<br>CTAACCATGTTATGCGGATCGCGCTTGAAGAACTGTTTGAGGCGATCCCTAATCTTGAGCGGGATACGCGCG<br>AGGGCGTTGAATTCTGGGGTTGGGGATTCCGTGGTCCGACTAGCTTGCACGTTACTTGGGAAGTT*TAATAG*G<br>GTACCAAGCTTCTCGAG |
| | **Amino Acid Sequence** | **MGGSSHHHHHHSSGENLYFQGH**MTTTERPDLAWLDEVTMTQLERNPYEVYERLRAEAPLAFVPVLGSYVAST<br>AEVCREVATSPDFEAVITPAGGRTFGHPAIIGVNGDIHADLRSMVEPALQPAEVDRWIDDLVRPIARRYLER<br>FENDGHAELVAQYCEPVSVRSLGDLLGLQEVDSDKLREWFAKLNRSFTNAAVDENGEFANPEGFAEGDQAKA<br>EIRAVVDPLIDRWIEHPDDSAISHWLHDGMPPGQTRDREYIYPTIYVYLLGAMETPGHGMASTLVGLFSRPE<br>QLEEVVDDPTLIPRAIAEGLRWTSPIWSATARISTKPVTIAGVDLPAGTPVMLSYGSANHDTGKYEAPSQYD<br>LHRPPLPHLAFGAGNHACAGIYFANHVMRIALEELFEAIPNLERDTREGVEFWGWGFRGPTSLHVTWEV |
| **GcoB pET29b** | **DNA Sequence** | TTCCTTCATATGACATTTGCCGTCTCTGTTGGAGGTCGTCGTGTTGACTGCGAGCCAGGGCAAACACTGCTT<br>GAAGCATTCCTGCGCGGTGGAGTCTGGATGCCTAATAGCTGTAACCAGGGGACCTGTGGGACATGTAAACTG<br>CAGGTATTATCAGGTGAAGTGGATCACGGAGCCGCCCCAGAAGATACGTTGTCCGCGGAGGAGCGTGCTGCT<br>GGATTAGCTTTAGCGTGCCAGGCACGTCCATTGGCAGACACGGAAGTGCGTTCGACAGCCGATGCAGGGCGT<br>GTAACCCATCCCTTGCGCGACCTTACCGCAACAGTGCTGGAGGTAGCAGACATCGCTCGTGACACTCGTCGC<br>GTCCTTTTAGGCTTGGCCGAGCCCTTGGCCTTTGAAGCGGGTCAATATGTGGAGTTGGTTGTTCCTGGGTCC<br>GGTGCGCGTCGCCAATACTCTTTAGCAAATACCGCAGACGAAGATAAGGTGCTTGAGTTGCATGTACGCCGC |

| | | |
|---|---|---|
| | | GTACCTGGGGGCATCGCCACAGATGGGTGGTTGTTTGATGGCCTGGCAGCAGGGGATCGTGTGGAAGCTAGT<br>GGACCCCTGGGTGACTTTCGTTTGCCACCACCTGAAGAGGACGATGGTGGACCAATGGTGTTGATCGGCGGG<br>GGGACAGGCCTGGCCCCCCTTGTCGGAATTGCCCGCACCGCATTGGCTCGTCATCCCTCTCGTGAGGTACTG<br>TTATATCACGGAGTCCGTGGGGTAGCAGATTTGTACGATTTGGGTCGTTTCGCTGAGATCGCGGAACAGCAT<br>CCGGGGTTTCGTTTCGTTCCCGTGCTGTCCGACGAACCGGATCCGGCTTATCGTTCAGGATTCCCAACTGAC<br>GCTTTTGTAGAAGACGTGCCCAGCGGGCGTGGTTGGTCGGGTTGGCTGTGCGGTCCCCCGGCGATGGTCGAA<br>GCAGGCGTTAAGGCTTTCAAACGCCGCCGCATGTCCCCACGTCGTATCCATCGTGAAAAGTTCACTCCTGCT<br>TCG*TAATAG*GGTACCAAGCTTTTAATT |
| | **Amino Acid Sequence** | MTFAVSVGGRRVDCEPGQTLLEAFLRGGVWMPNSCNQGTCGTCKLQVLSGEVDHGAAPEDTLSAEERAAGLA<br>LACQARPLADTEVRSTADAGRVTHPLRDLTATVLEVADIARDTRRVLLGLAEPLAFEAGQYVELVVPGSGAR<br>RQYSLANTADEDKVLELHVRRVPGGIATDGWLFDGLAAGDRVEASGPLGDFRLPPPEEDDGGPMVLIGGGTG<br>LAPLVGIARTALARHPSREVLLYHGVRGVADLYDLGRFAEIAEQHPGFRFVPVLSDEPDPAYRSGFPTDAFV<br>EDVPSGRGWSGWLCGPPAMVEAGVKAFKRRRMSPRRIHREKFTPAS |
| **GcoB pET29b** | **DNA Sequence** | CATATGACATTTGCCGTCTCTGTTGGAGGTCGTCGTGTTGACTGCGAGCCAGGGCAAACACTGCTTGAAGCA<br>TTCCTGCGCGGTGGAGTCTGGATGCCTAATAGCTGTAACCAGGGGACCTGTGGGACATGTAAACTGCAGGTA<br>TTATCAGGTGAAGTGGATCACGGAGCCGCCCCAGAAGATACGTTGTCCGCGGAGGAGCGTGCTGCTGGATTA<br>GCTTTAGCGTGCCAGGCACGTCCATTGGCAGACACGGAAGTGCGTTCGACAGCCGATGCAGGGCGTGTAACC<br>CATCCCTTGCGCGACCTTACCGCAACAGTGCTGGAGGTAGCAGACATCGCTCGTGACACTCGTCGCGTCCTT<br>TTAGGCTTGGCCGAGCCCTTGGCCTTTGAAGCGGGTCAATATGTGGAGTTGGTTGTTCCTGGGTCCGGTGCG<br>CGTCGCCAATACTCTTTAGCAAATACCGCAGACGAAGATAAGGTGCTTGAGTTGCATGTACGCCGCGTACCT<br>GGGGGCATCGCCACAGATGGGTGGTTGTTTGATGGCCTGGCAGCAGGGGATCGTGTGGAAGCTAGTGGACCC<br>CTGGGTGACTTTCGTTTGCCACCACCTGAAGAGGACGATGGTGGACCAATGGTGTTGATCGGCGGGGGACA<br>GGCCTGGCCCCCCTTGTCGGAATTGCCCGCACCGCATTGGCTCGTCATCCCTCTCGTGAGGTACTGTTATAT<br>CACGGAGTCCGTGGGGTAGCAGATTTGTACGATTTGGGTCGTTTCGCTGAGATCGCGGAACAGCATCCGGGG<br>TTTCGTTTCGTTCCCGTGCTGTCCGACGAACCGGATCCGGCTTATCGTTCAGGATTCCCAACTGACGCTTTT<br>GTAGAAGACGTGCCCAGCGGGCGTGGTTGGTCGGGTTGGCTGTGCGGTCCCCCGGCGATGGTCGAAGCAGGC<br>GTTAAGGCTTTCAAACGCCGCCGCATGTCCCCACGTCGTATCCATCGTGAAAAGTTCACTCCTGCTTCG**CAC**<br>**CACCACCACCACCAC***TAATAG*AAGCTT |
| | **Amino Acid Sequence** | MTFAVSVGGRRVDCEPGQTLLEAFLRGGVWMPNSCNQGTCGTCKLQVLSGEVDHGAAPEDTLSAEERAAGLA<br>LACQARPLADTEVRSTADAGRVTHPLRDLTATVLEVADIARDTRRVLLGLAEPLAFEAGQYVELVVPGSGAR<br>RQYSLANTADEDKVLELHVRRVPGGIATDGWLFDGLAAGDRVEASGPLGDFRLPPPEEDDGGPMVLIGGGTG<br>LAPLVGIARTALARHPSREVLLYHGVRGVADLYDLGRFAEIAEQHPGFRFVPVLSDEPDPAYRSGFPTDAFV<br>EDVPSGRGWSGWLCGPPAMVEAGVKAFKRRRMSPRRIHREKFTPAS**HHHHHH** |

**Table S6.** Sequences of DNA primers used in this study.

| Primer ID | Direction | Sequence (5'–3') |
|---|---|---|
| **215** | FWD | TAATACGACTCACTATAGG |
| **315** | REV | GCTAGTTATTGCTCAGCGGTGG |
| **1330** | REV | GTGAAATACCGCACAGATGCGTAAG |
| **3497** | FWD | TGAGGATCCGAATTCGAGCTCC |
| **2281** | REV | GCCCTGGAAGTACAGGTTTTCG |
| **3312** | FWD | TTCTCCTTACGCATCTGTGCGG |
| **3558** | FWD | ACCACCGAAAACCTGTACTTCCAGGGCGCCATGACAACAAAGCATACTACAGCCGG |
| **3559** | REV | AGCTTGTCGACGGAGCTCGAATTCGGATCCTCAGTCCCACAGCACATCCAAAC |
| **RS0114865 5'** | FWD | TTAATTCATATGACGCGTGAGCTGGTGG |
| **RS0114865 His 3'** | REV | TTAATTAAGCTTCTATTAGTGGTGGTGGTGGTGGTGAAGATCCAACCCCAGCACCGCTG |
| **RS0109265 5'** | FWD | TTAATTCATATGACATTTGCCGTCTCTG |
| **RS0109265 His 3'** | REV | TTAATTAAGCTTCTATTAGTGGTGGTGGTGGTGGTGCGAAGCAGGAGTGAACTTTTCAC |
